# Single-cell mapping reveals new markers and functions of lymphatic endothelial cells in lymph nodes

**DOI:** 10.1101/2020.01.09.900241

**Authors:** Noriki Fujimoto, Yuliang He, Marco D’Addio, Carlotta Tacconi, Michael Detmar, Lothar C. Dieterich

**Affiliations:** Institute of Pharmaceutical Sciences, Swiss Federal Institute of Technology (ETH) Zurich, Zurich, Switzerland; Department of Dermatology, Shiga University of Medical Sciences, Japan

## Abstract

Lymph nodes (LNs) are highly organized secondary lymphoid organs that mediate adaptive immune responses to antigens delivered via afferent lymphatic vessels. Lymphatic endothelial cells (LECs) line intranodal lymphatic sinuses and organize lymph and antigen distribution. LECs also directly regulate T cells, mediating peripheral tolerance to self-antigens, and play a major role in many diseases including cancer metastasis. However, little is known about the phenotypic and functional heterogeneity of LN LECs. Using single-cell RNA sequencing, we comprehensively defined the transcriptome of LECs in murine skin-draining LNs, and identified new markers and functions of distinct LEC subpopulations. We found that LECs residing in the subcapsular sinus have an unanticipated function in scavenging of modified LDL and also identified a specific cortical LEC subtype implicated in rapid lymphocyte egress from LNs. Our data provide new insights into the diversity of LECs in murine lymph nodes and a rich resource for future studies into the regulation of immune responses by lymph node LECs.

## INTRODUCTION

Peripheral lymph nodes (LNs) are essential secondary lymphoid organs that mediate interactions between antigen-presenting cells and lymphocytes for the initiation of adaptive immune responses. LNs also act as filters that retain specific proteins and other biomolecules present in the afferent lymph (Clement et al., 2018). Apart from lymphocytes, LNs comprise several stromal cell types, including fibroblastic reticular cells (FRCs), blood vascular endothelial cells, and lymphatic endothelial cells (LECs), that are crucial for LN development and function. LECs do not only provide structure to the LN sinuses that allow lymph percolation through the node, but also control the access of soluble molecules and subcellular particles (including viruses) to the conduit system that guides them to dendritic cells residing in the LN cortex (Rantakari et al., 2015; Reynoso et al., 2019). LN LECs also actively engage in a variety of immune-related processes. Under steady-state conditions, LN LECs control lymphocyte egress from LNs via generation of an S1P gradient (Cyster and Schwab, 2012) and regulate peripheral tolerance by expression and presentation of peripheral tissue self-antigens in combination with constitutive expression of PD-L1 and other regulatory molecules, leading to inhibition or deletion of auto-reactive CD8^+^ T cells (Cohen et al., 2010; Tewalt et al., 2012). Furthermore, similar to antigen-presenting cells, LN LECs have been reported to scavenge and (cross-)present exogenous antigen taken up from the lymph (Hirosue et al., 2014).

LNs draining inflamed tissues or tumors commonly increase in size, which is accompanied by an expansion of LECs (Dieterich and Detmar, 2016; Dieterich et al., 2014). LN lymphangiogenesis may be driven by soluble factors drained from the upstream tissue, or by signals produced locally in the lymph node, such as B cell-derived VEGF-A (Angeli et al., 2006). Consequently, in the case of tumor-draining LNs, lymphatic expansion can even occur before the colonization by tumor cells (Hirakawa et al., 2007), a process that may be involved in the generation of a “pre-metastatic niche” (Karaman and Detmar, 2014). Importantly, in pathological conditions LN LECs do not only increase in number, but also adapt a distinctive molecular phenotype. Several studies have characterized transcriptional changes of bulk-isolated LN LECs in response to experimental inflammation, virus infection and in the context of upstream tumor growth (Commerford et al., 2018; Gregory et al., 2017; Malhotra et al., 2012), demonstrating that LN LECs dynamically regulate a large number of genes associated with inflammatory processes, which conceivably affects their function and subsequently, any adaptive immune responses generated in the node.

LNs are highly organized structures that host specialized immune cell types in defined anatomical compartments, such as subcapsular, cortical and medullary regions. Therefore, it is conceivable that stromal cells parallel this zonation and display diverse phenotypes and functions, depending on their location in the node. For example, Rodda et al. recently identified multiple subtypes of FRCs that differed in location and gene expression (Rodda et al., 2018). Several reports have addressed the heterogeneity of LECs in murine LNs, typically focusing on selected marker genes only. One of the most prominent examples are the LECs lining the ceiling and the floor of the subcapsular sinus (SCS), which exhibit markedly different phenotypes, in spite of their close physical proximity. For example, ceiling LECs (cLECs) express the atypical chemokine receptor ACKR4 and display no or only low levels of LYVE1, whereas floor-lining LECs (fLECs) express high levels of LYVE1 but are negative for ACKR4 (Ulvmar et al., 2014). Further studies identified MADCAM1 and ITGA2B as additional markers of fLECs (Bovay et al., 2018; Cohen et al., 2014; Cordeiro et al., 2016). The distinct molecular phenotypes of c- and fLECs likely enable them to support specific functions, such as the fLEC-specific transmigration of antigen-presenting cells (Braun et al., 2011; Ulvmar et al., 2014) or transcytosis of antibodies (Kahari et al., 2019) from the subcapsular sinus into the LN cortex. However, to comprehensively define the molecular and functional heterogeneity of LN LECs, analysis at single cell resolution is necessary.

Recently, a study by Takeda et al. reported a single-cell sequencing analysis of LN LECs isolated from cancer patients, and described four subsets corresponding to subcapsular sinus cLECs and fLECs, a second type of cLECs present only in the medullary region, and a single cluster of medullary and cortical LECs which were transcriptionally indistinguishable (Takeda et al., 2019). However, sequencing depth of the transcriptional data provided in this study was relatively shallow, and might have been influenced by the diseased state due to systemic responses to tumor growth. Here, we performed single-cell RNA sequencing of LECs isolated from murine inguinal LNs of completely naïve animals for unbiased identification of LEC subsets and comprehensive characterization of their phenotypes in steady state. Our results reveal that there are at least four subsets of murine LN LECs with marked differences in gene expression that correspond to distinct anatomical locations within the LN. These included the cLECs and fLECs of the SCS, as well as medullary sinus LECs, similar to what has been described in humans (Takeda et al., 2019). Notably, we additionally identified a small subset of cortical sinuses that mediate rapid lymphocyte egress from the LN, and we uncovered a hitherto unknown function of LN LECs in scavenging low-density lipoprotein (LDL).

## RESULTS

### Identification of four LN LEC subtypes and gene expression signatures by single cell sequencing

To map the heterogeneity of LN LECs, we isolated these cells (CD45^−^CD31^+^podoplanin^+^, Figure 1A) from inguinal LNs of C57Bl/6 wildtype mice by FACS sorting and subjected them to deep RNA sequencing at single cell resolution (N=1152 cells) using the SmartSeq2 full-length transcriptome profiling approach (Picelli et al., 2014). After quality filtering to remove cells with outlier read counts (N=134) and a group of cells showing CD45 expression (N=125, data not shown) which were probably due to sorting impurity, 893 cells were subjected to further analysis. Unsupervised clustering suggested the existence of at least 4 LEC subtypes: the largest cluster (cluster 3) comprised 364 cells (40.8%), cluster 1 283 cells (31.7%), cluster 2 194 cells (21.7%), and the smallest cluster (cluster 4), located between cluster 2 and 3, 52 cells (5.8%) (Figure 1B). All of these cells showed robust expression of the two markers used for FACS sorting, CD31 (Pecam1) and podoplanin (Pdpn) (Figure 1C), the pan-endothelial marker VE-cadherin (Cdh5, data not shown), and the LEC marker genes Prox1 and Flt4 (Vegfr3) (Figure 1C), confirming their lymphatic endothelial identity. Previously, it has been reported that LECs lining the subcapsular sinus show distinct marker expression depending on their location in the ceiling or the floor of the sinus. For example, cLECs are negative for LYVE1 and ITGA2B, but express the atypical chemokine receptor ACKR4, whereas fLECs express MADCAM1 (Bovay et al., 2018; Cordeiro et al., 2016; Ulvmar et al., 2014) (Figure S1A). In agreement with this, we observed differential expression of these genes among the four LEC clusters. LYVE1 and ITGA2B were present in all clusters apart from cluster 2; cluster 1 specifically expressed MADCAM1; and cluster 2 specifically expressed ACKR4 (Figure 1D). This suggests that the clusters we identified based on gene expression correspond to LECs in different anatomical locations in the LN, with cluster 1 representing fLECs and cluster 2 cLECs, whereas clusters 3 and 4 most likely represent cortical and / or medullary LEC subsets.

**Figure 1.**
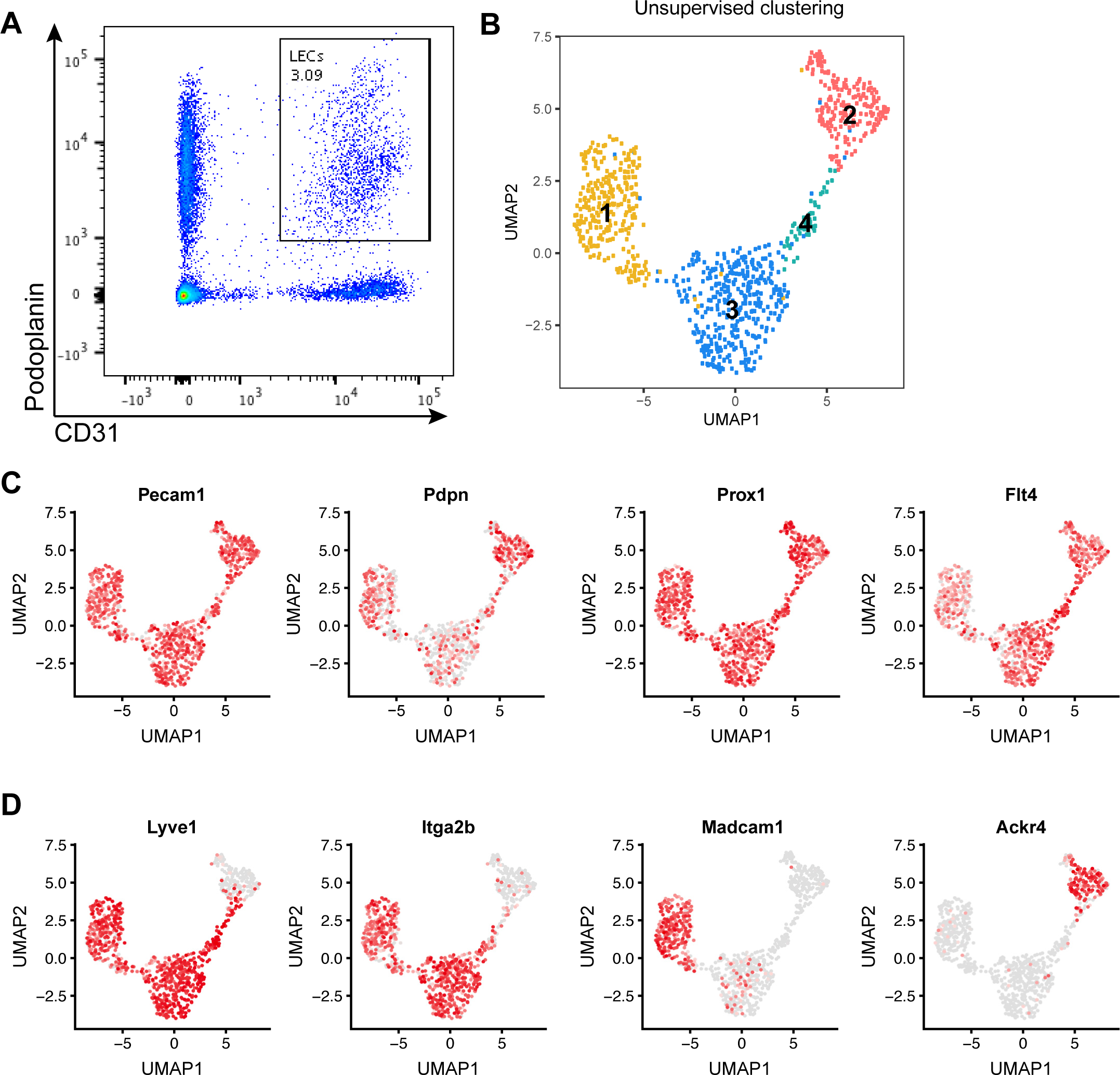
Single cell transcriptomic analysis of LN LECs. (A) Example FACS plot of LN stromal cells (pregated as CD45^−^ living singlets) from inguinal LNs of C57Bl/6 wildtype mice. CD31^+^ podoplanin^+^ LECs were isolated by single-cell sorting. The number in the LEC gate indicates the percentage of LECs among all stromal cells. (B) Unsupervised clustering of 893 LN LECs resulted in 4 distinct clusters. Each point represents an individual cell. (C, D) Expression levels of selected genes plotted using the original log-transformed counts. Grey dots indicate cells without any measurable expression; red dots coded by color intensity denote the detected expression magnitude. (C) Pecam1 (CD31) and Pdpn used as markers for FACS sorting as well as the LEC marker genes Prox1 and Flt4 (Vegfr3) were robustly expressed in most cells. (D) While Lyve1 and Itga2b were expressed in all clusters except for cluster 2, the fLEC marker Madcam1 and the cLEC marker Ackr4 were specifically expressed in cluster 1 and cluster 2, respectively.

Differential expression (DE) analysis among the 4 clusters identified significantly (log_2_ FC ≥ 0.6; FDR < 0.01) up- or downregulated genes in each of the clusters compared to all others. Interestingly, gene ontology (GO) analysis of these DE genes suggested “opposing” phenotypes of cluster 1 LECs (fLECs) and cluster 2 LECs (cLECs), with an enrichment of inflammation-associated genes and a de-enrichment of angiogenesis-associated transcripts in cluster 1, and vice versa an enrichment of angiogenesis-associated and a de-enrichment of inflammation-associated genes in cluster 2 (Supplementary Table 1). Cluster 4 LECs also showed an enrichment of angiogenesis-related transcripts, whereas cluster 3 was characterized by metabolism- and oxidation-related terms (Supplementary Table 1).

### Molecular characterization of LECs in the SCS floor

To confirm the identity and the anatomical location of the LEC clusters, we performed immunofluorescence staining and RNA detection *in situ* in inguinal LN tissue sections. To evaluate the distribution of marker expression, we selected three regions within the LN: (1) the SCS associated with B cell follicles; (2) the SCS and large interfollicular sinus tracts entering the nodes between the follicles (denoted as IF-SCS); (3) and the medullary sinuses within the node (MS) (Figure 2A). To validate that cluster 1 LECs were accurately assigned to the fLECs, we first analyzed the expression of the known fLEC-expressed genes ITGA2B and LYVE1. As expected, these two markers were clearly detectable in fLECs in the SCS and more broadly in the IF-SCS regions (Figure 2B, S1B). Similarly, ACKR4^+^ cLECs were detectable in both the SCS and the IF-SCS region (Figure S1C-D).

**Figure 2.**
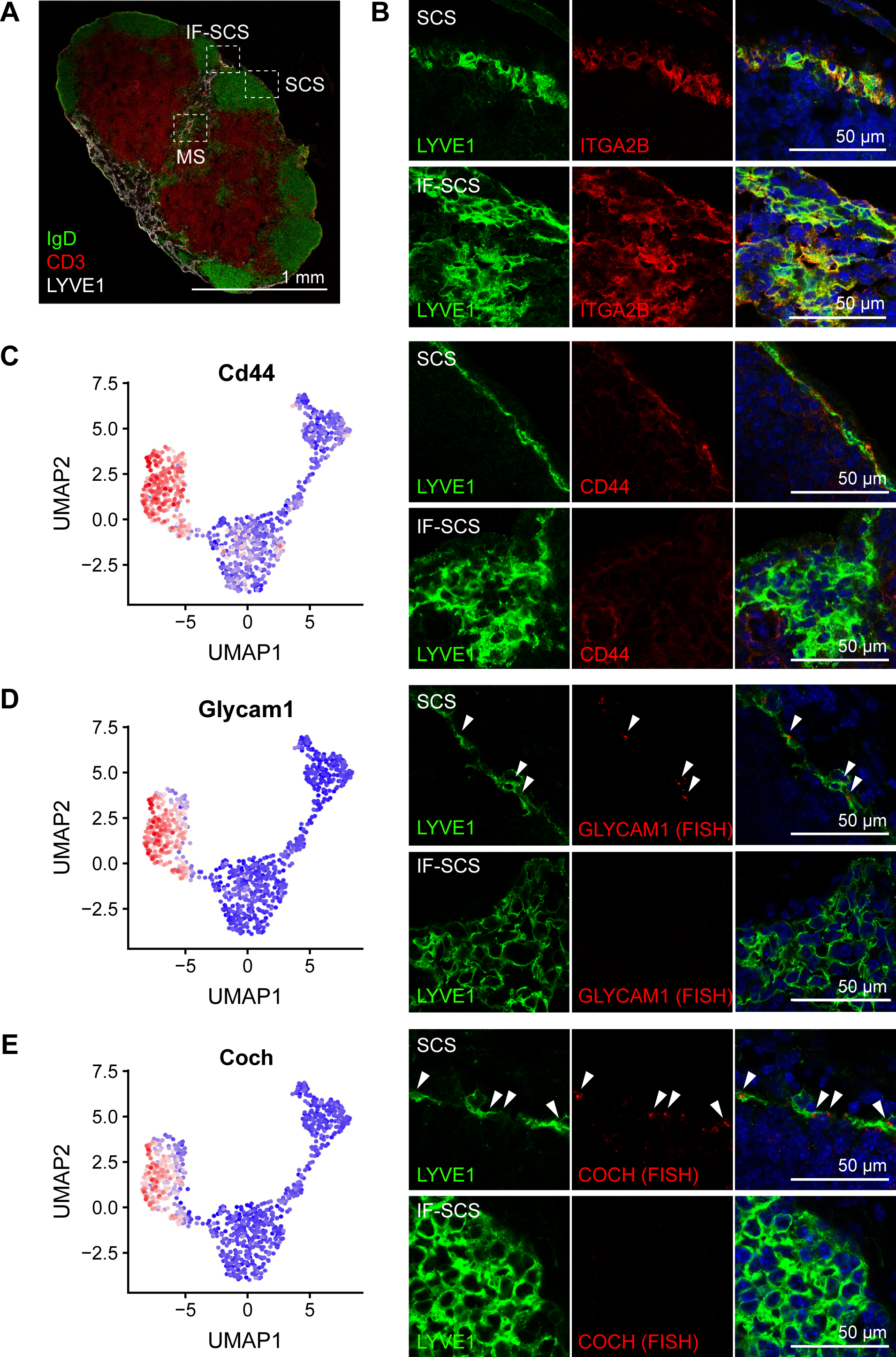
Molecular characterization of LECs in the SCS floor. (A) Immunofluorescence staining of a whole inguinal LN cross-section for IgD (green), CD3 (red) and LYVE1 (white). The dotted squares indicate three regions that were used for further analysis: the SCS (subcapsular sinus associated with B cell follicles), the IF-SCS (SCS and large interfollicular sinus tracts entering the node), and the MS (medullary sinuses within the node). (B) Immunofluorescence co-staining for LYVE1 (green) and ITGA2B (red) in the SCS (upper panels) and the IF-SCS region (lower panels) showed expression of ITGA2B and LYVE1 by LECs in both the SCS floor and the IF-SCS region. (C-E) Gene expression patterns (left panels; denoted as high expression level in red and low in blue, using the corrected expression values) and protein / transcript location of new fLEC (cluster 1) markers. CD44 (C, detected by immunofluorescence staining), Glycam1 (D, detected by RNA FISH), and Coch (E, detected by RNA FISH) were specifically expressed in fLECs in the SCS region (upper right panels), but not in the IF-SCS region (lower right panels). White arrowheads indicate RNA FISH signals; LYVE1 stained in green.

Next, we chose several new markers significantly upregulated in cluster 1 / fLECs for validation. CD44, a homologue of LYVE1 (Banerji et al., 1999), was specifically expressed in cluster 1 LECs. In agreement with this, immunofluorescence staining of CD44 was found in LYVE1^+^ fLECs in the SCS region. Interestingly however, it was largely absent from the IF-SCS region (Figure 2C), suggesting that the cluster 1 / fLEC subset is only present in the SCS right above B cell follicles, whereas most of the LECs in IF-SCS regions display a different phenotype that rather corresponds to cluster 3. In line with this, Glycam1 and cochlin (coch), two additional cluster 1-specific transcripts, could also be detected in fLECs in the SCS region by *in situ* RNA hybridization but were absent from the IF-SCS region (Figure 2D-E).

### Identification of new markers and functions of LECs in the SCS ceiling

Since ACKR4 is a well-established marker of cLECs (Ulvmar et al., 2014), we used Ackr4-GFP reporter mice to characterize the expression of potential new cLEC marker genes. Of note, cLECs (cluster 2 LECs) were the most distinguishable LN LEC subset in our dataset, with a total of 220 up- and 149 downregulated genes in this cluster compared to all the other clusters. We selected several of these genes that were suitable for immunofluorescence staining, and investigated their expression patterns in inguinal LNs. ANXA2 was highly expressed in cluster 2 / cLECs in our dataset, and we correspondingly found it located in the SCS ceiling and the capsule, partially overlapping with ACKR4-driven GFP expression (Figure 3A). Interestingly, ANXA2 also stained afferent lymphatic vessels merging with the SCS, demonstrating a phenotypic similarity between cLECs and afferent lymphatic collectors (Figure S2A). In addition, many immune cells, particularly within the T cell zone of the node, stained for ANXA2 (data not shown). Similarly, FABP4, CD36, FLRT2 and BGN were also confined to the SCS ceiling (Figure 3B-C and S2B-C). Our sequencing data furthermore indicated that cLECs specifically express another atypical chemokine receptor, Ackr3, as well as Btnl9 which is related to the co-stimulatory B7 gene (Abeler-Dorner et al., 2012). Owing to the shortage of commercially available antibodies, we localized the corresponding transcripts using RNA *in situ* hybridization. Expression of both Ackr3 and Btnl9 mRNA was detectable in various regions and cell types of inguinal LNs, but within the lymphatic endothelium, was confined to cLECs (Figure 4A-B).

**Figure 3.**
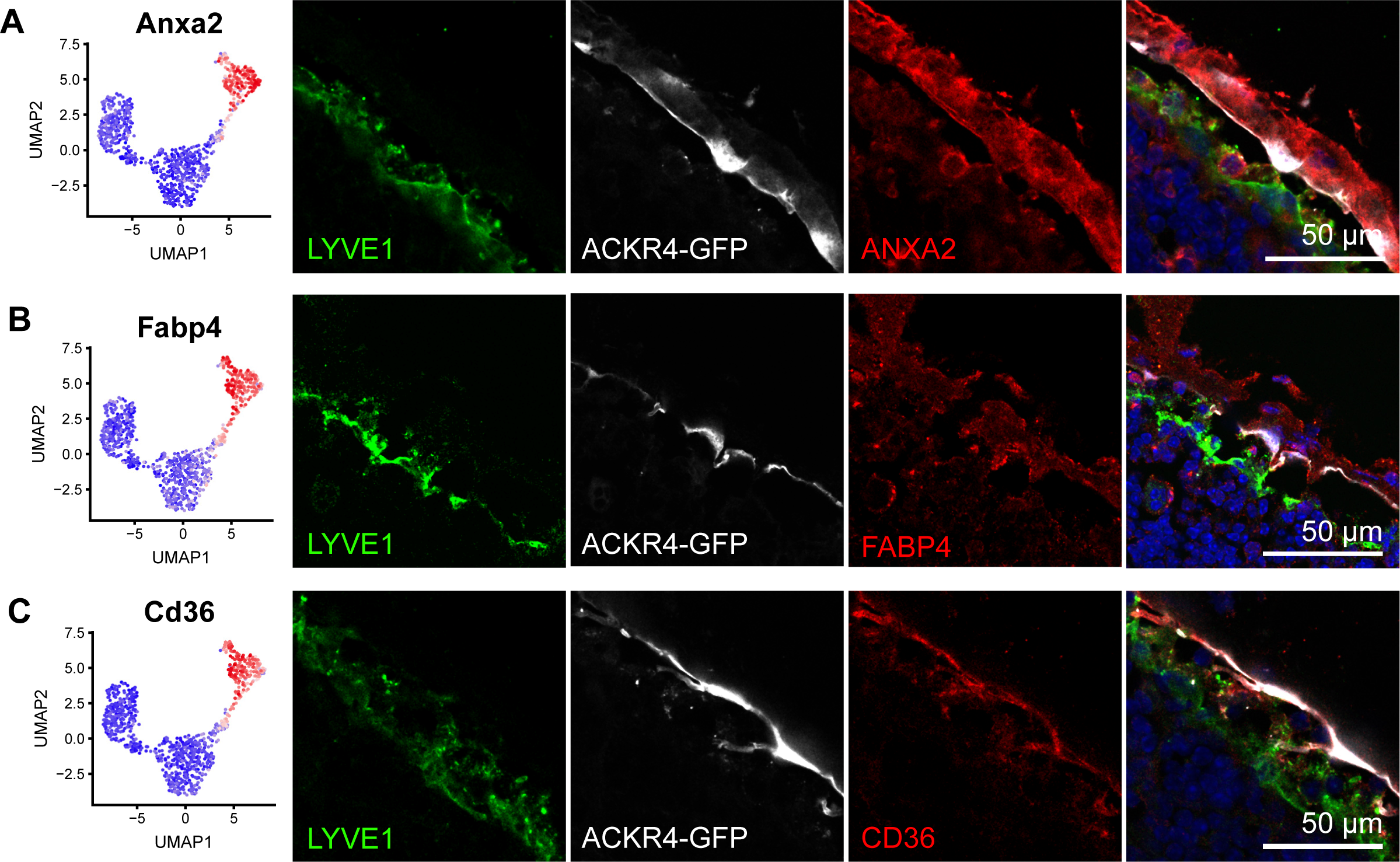
Molecular characterization of LECs in the SCS ceiling with immunofluorescence staining. (A-C) Expression of new cLEC / cluster 2 marker genes ANXA2 (A), FABP4 (B) and CD36 (C) by RNA sequencing (left panels) and immunofluorescence staining (right panels) in Ackr4-GFP reporter mice. GFP (white) and immunofluorescence co-staining for LYVE1 (green) served as markers for cLECs and fLECs, respectively.

**Figure 4.**
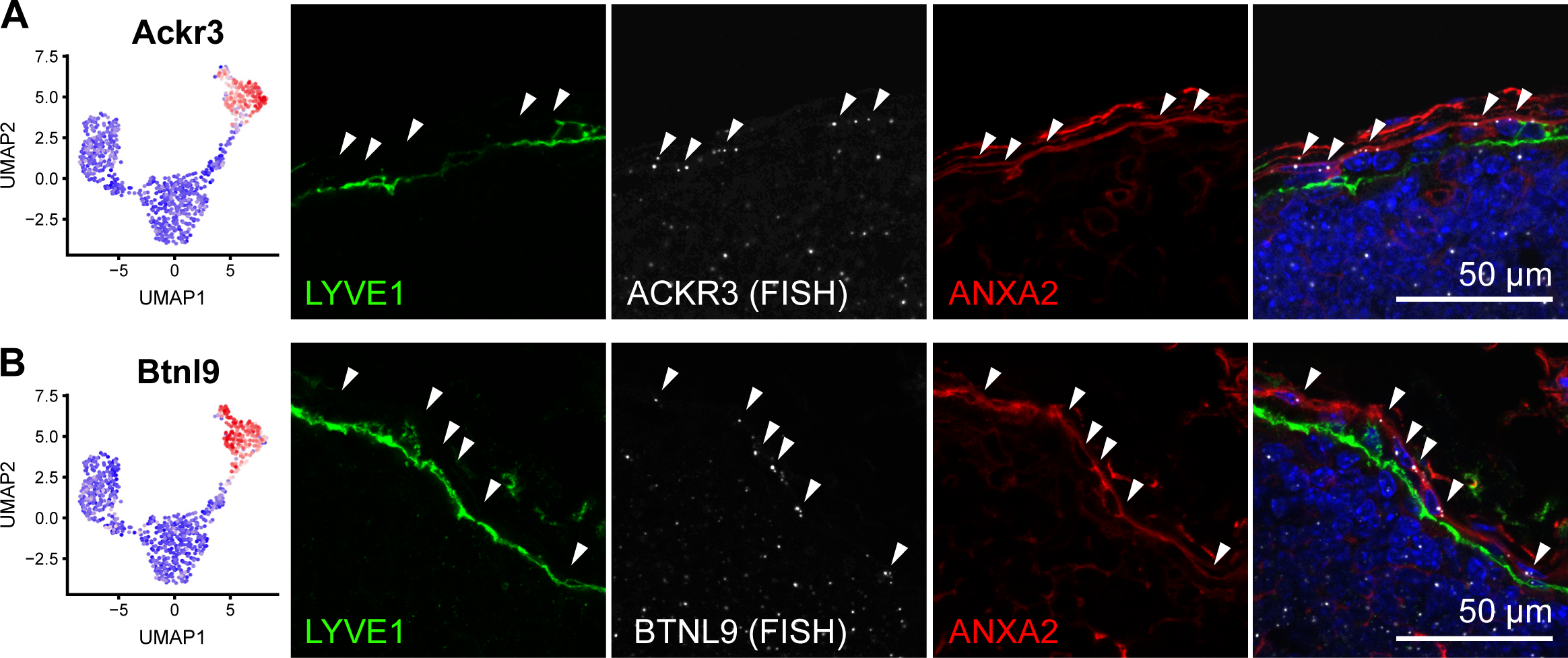
Molecular characterization of LECs in the SCS ceiling with RNA FISH. (A-B) Expression of new cLEC / cluster 2 marker genes Ackr3 (A) and Btnl9 (B) by RNA sequencing (left panels) and RNA FISH (right panels). As GFP fluorescence is lost during tissue processing for RNA FISH, immunofluorescence staining for ANXA2 (red) and LYVE1 (green) served as markers for cLECs and fLECs, respectively. Arrows point to cLECs expressing Ackr3 and Btnl9 transcripts (white).

Surprisingly, differential expression analysis identified genes known to be involved in the uptake of modified LDLs (Levitan et al., 2010). For example, CD36 was specifically expressed in cLECs (Figure 3), whereas Msr1 and Fcgr2b were excluded from cLECs but highly expressed in most other LN LECs. This prompted us to evaluate whether cLECs would have a distinct capacity to take up modified LDL from the lymph. To this end, we injected Ackr4-GFP mice intradermally with fluorescently-labeled acetylated or oxidized LDL near the base of the tail and collected the draining inguinal LNs 1 h later. Histological analysis revealed striking differences in LDL distribution in the LN LECs. Acetylated LDL partly overlapped with ACKR4^+^ cLECs, indicating selective uptake by this LEC subset (Figure 5A-B), whereas oxidized LDL was rather taken up by LYVE1^+^ LECs in the SCS floor and in cortical regions (Figure 5C-D). These data reveal a novel function of LECs in scavenging of LDLs from the lymph, and furthermore suggest that cLECs and other LN LECs have a distinct capacity to take up differentially modified LDL.

**Figure 5.**
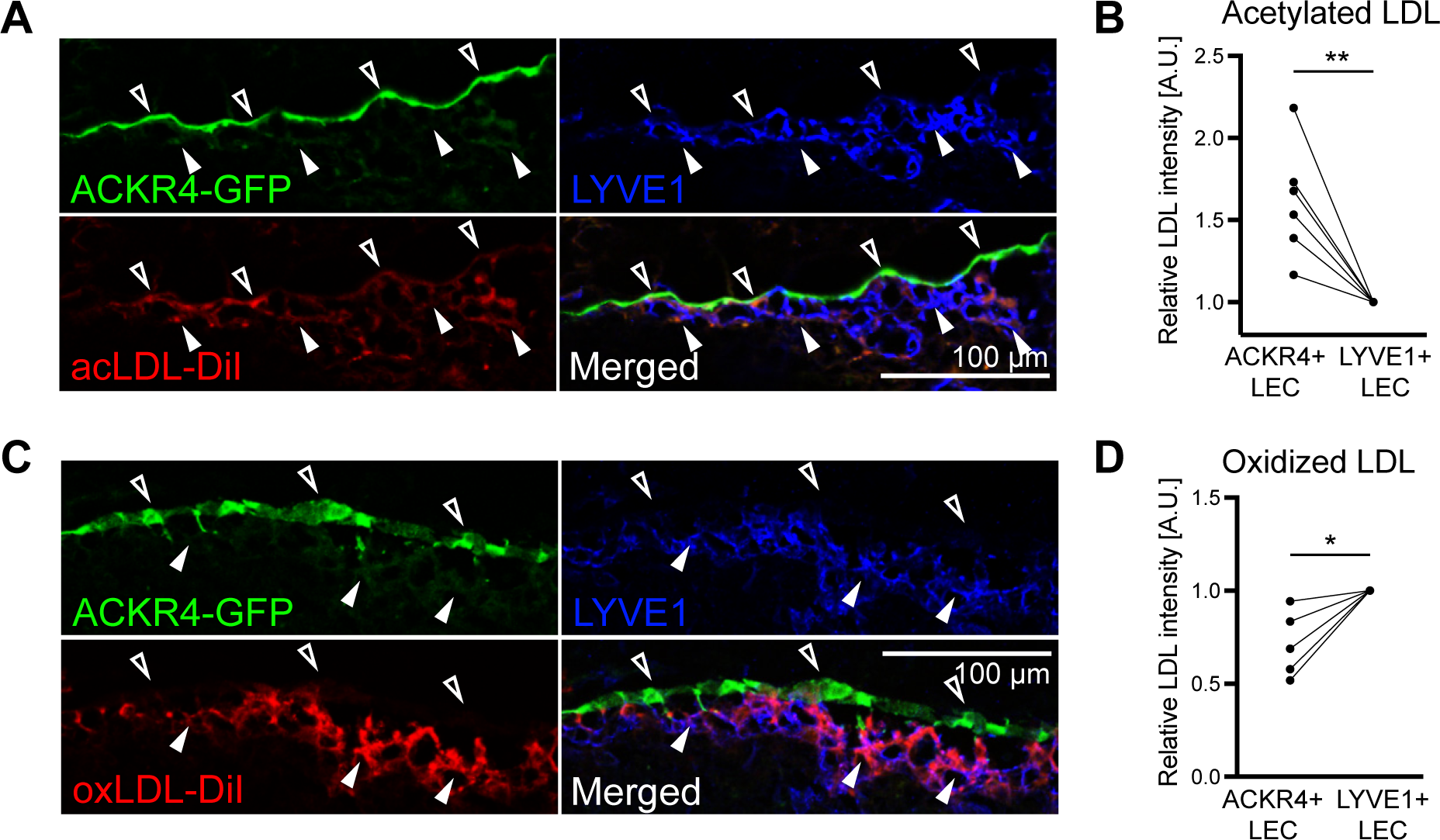
Differential LDL uptake by cLECs. *In vivo* LDL tracing after intradermal injection of Dil-labeled acetylated or oxidized LDL near the base of the tail of Ackr4-GFP reporter mice. Draining inguinal LNs were collected 1 h later. (A, B) Representative images (A) and quantification (B) of acetylated LDL (red) accumulation in ACKR4+ and LYVE1+ LECs. Each line represents one mouse (n = 6). (C-D) Representative images (C) and quantification (D) of oxidized LDL (red) accumulation in ACKR4+ and LYVE1+ LECs. Filled arrowheads indicate cLECs, empty arrowheads indicated fLECs. For quantification, signal intensities were normalized to the level of ACKR4-LYVE1+ LECs (n = 5). ** p < 0.01; * p < 0.05 (paired t-test).

### LECs in medullary and interfollicular sinuses are phenotypically similar

Cluster 3 was the most abundant LEC subset in our sequencing dataset, and therefore likely represented cells lining the medullary and / or cortical sinuses. To test this hypothesis, we chose several marker genes expressed by cluster 3, namely IL33 (which was also expressed in fLECs) as well as MRC1 and MARCO, two genes typically associated with macrophages but that have previously been shown to be expressed by human LN LECs (Martens et al., 2006). Immunofluorescence staining confirmed expression of IL33 in fLECs in the SCS, and most LECs in the IF-SCS and MS regions (Figure 6A). Conversely, MRC1 and MARCO were absent from fLECs as expected, but were expressed by medullary LECs, both in IF-SCS and MS regions (Figure 6B-C). These data indicate that cluster 3 LECs represent large interfollicular and medullary sinuses, which consequently appear to have a very similar phenotype.

**Figure 6.**
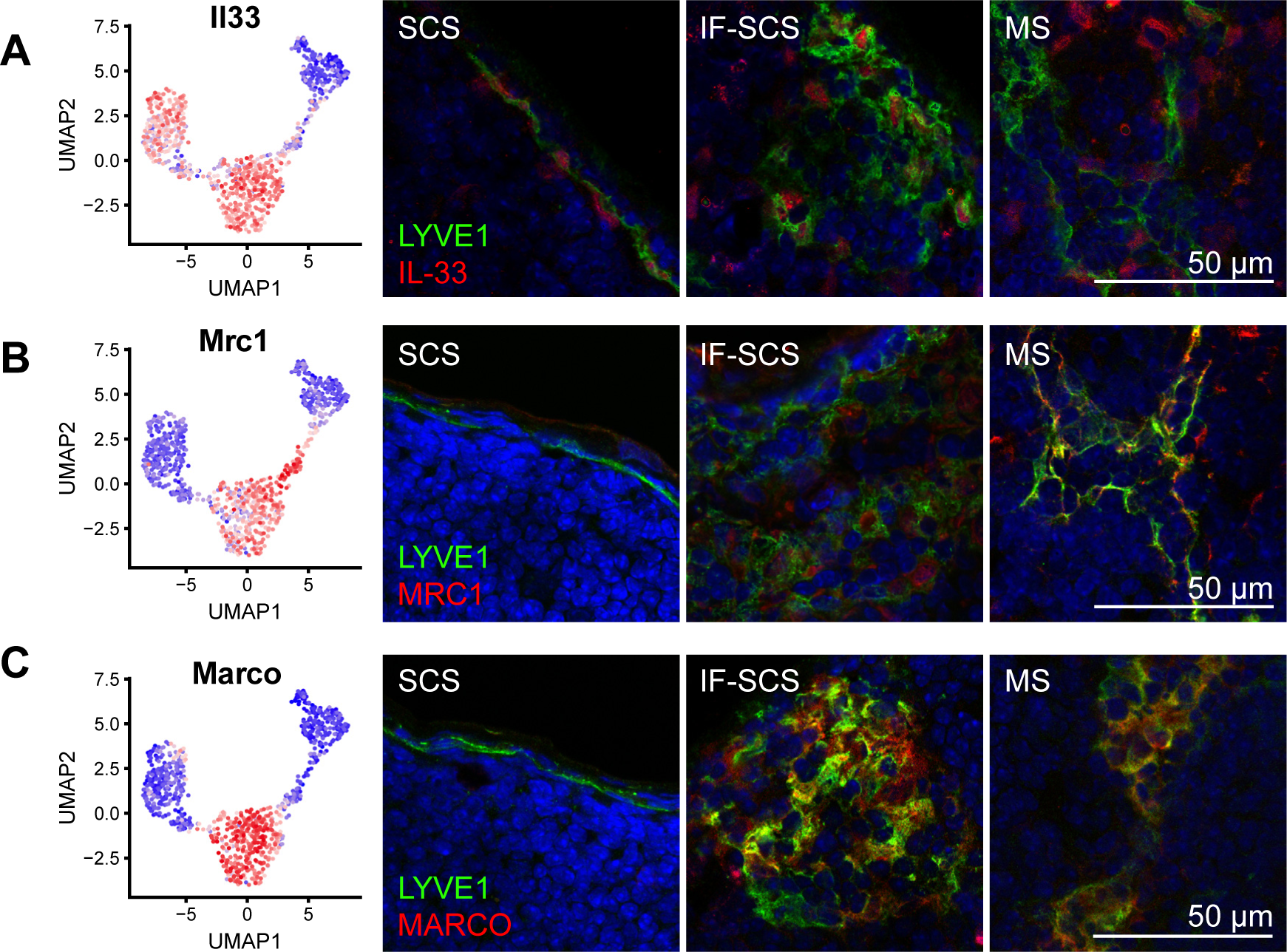
Molecular characterization of medullary sinus LECs. (A-C) Expression of MS LEC / cluster 3 marker genes IL33 (A), MRC1 (B) and MARCO (C) by RNA sequencing (left panels) and immunofluorescence staining (right panels, red) in combination with LYVE1 (green). IL33 (A) was expressed by fLECs and MS LECs and showed nuclear LEC staining in the SCS, IF-SCS and MS regions, whereas MRC1 (B) and MARCO (C) were excluded from fLECs and congruously showed no staining in the SCS region.

### New LEC subset-specific markers are largely conserved among LNs from various anatomical locations and allow subset discrimination by flow cytometry

Next, we sought to investigate whether the expression pattern of the new markers for cLECs, fLECs and medullary LECs we identified in inguinal LNs would be similar in LNs residing at other anatomical locations and therefore draining other organs than the skin, such as mandibular (draining facial regions as well as the brain (Ma et al., 2017)), iliac (draining predominantly the lower gastrointestinal tract) and mesenteric LNs (draining the upper gastrointestinal tract). Immunofluorescence staining of several selected markers (CD44, ANXA2, CD36, MRC1) revealed a remarkable conservation in these nodes, with the sole exception of CD44 which was not expressed in mesenteric fLECs (Figure S3). Additionally, we found that some of these markers are also suitable to discriminate between the major LN LEC subsets by flow cytometry. Using inguinal LNs from Ackr4-GFP mice, a combination of ITGA2B, CD44 and MRC1 allowed us to distinguish between cLECs (GFP^+^, MRC1^−^, ITGA2B^−^, CD44^lo^), fLECs (GFP^−^, MRC1^−^, ITGA2B^+^, CD44^+^), and medullary LECs (GFP^−^, MRC1^+^, ITGA2B^+/lo^, CD44^−^) (Figure S4).

### A unique subset of cortical and medullary sinuses serves as lymphocyte egress structures

The smallest LN LEC subset, cluster 4, shared the expression of many genes with medullary LECs (cluster 3) and with cLECs (cluster 2). For example, cluster 4 LECs expressed both LYVE1 as well as intermediate levels of the otherwise cLEC-restricted marker ANXA2 (Figure 1D, 3A). Interestingly, immunofluorescence staining of these two markers identified a subset of lymphatic sinuses located in the (para-) cortex, close to the medulla of inguinal LNs, frequently in proximity to high endothelial venules (HEVs) that were strongly positive for Glycam1 (Imai et al., 1991) (Figure 7A). Furthermore, they were surrounded and filled by B and T lymphocytes, but rarely by F4/80^+^ or CD169^+^ macrophages, further distinguishing them from medullary sinuses (Figure S5A-D). We also confirmed that those structures were indeed lymphatic sinuses by staining for Prox1, and that they expressed MRC1 but were negative for MARCO (Figure S5E-G) as suggested by the RNA sequencing data (Figure 6B-C). To further confirm that cluster 4 LECs indeed correspond to those sinuses, we selected several transcripts specific for this cluster, namely Ptx3, Kcnj8, and Itih5, and mapped them by *in situ* RNA hybridization. In all cases, expression outside of lymphatic sinuses could be detected (data not shown), which might be derived from other LN stromal cells or immune cells. However, within LYVE1^+^ lymphatic structures, these transcripts were only detectable in ANXA2^+^ sinuses (Figure 7B-D). Taken together, this demonstrates that the cluster 4 LECs identified by single cell RNA sequencing correspond to a unique subset of lymphatic sinuses in the cortex of mouse inguinal LNs.

**Figure 7.**
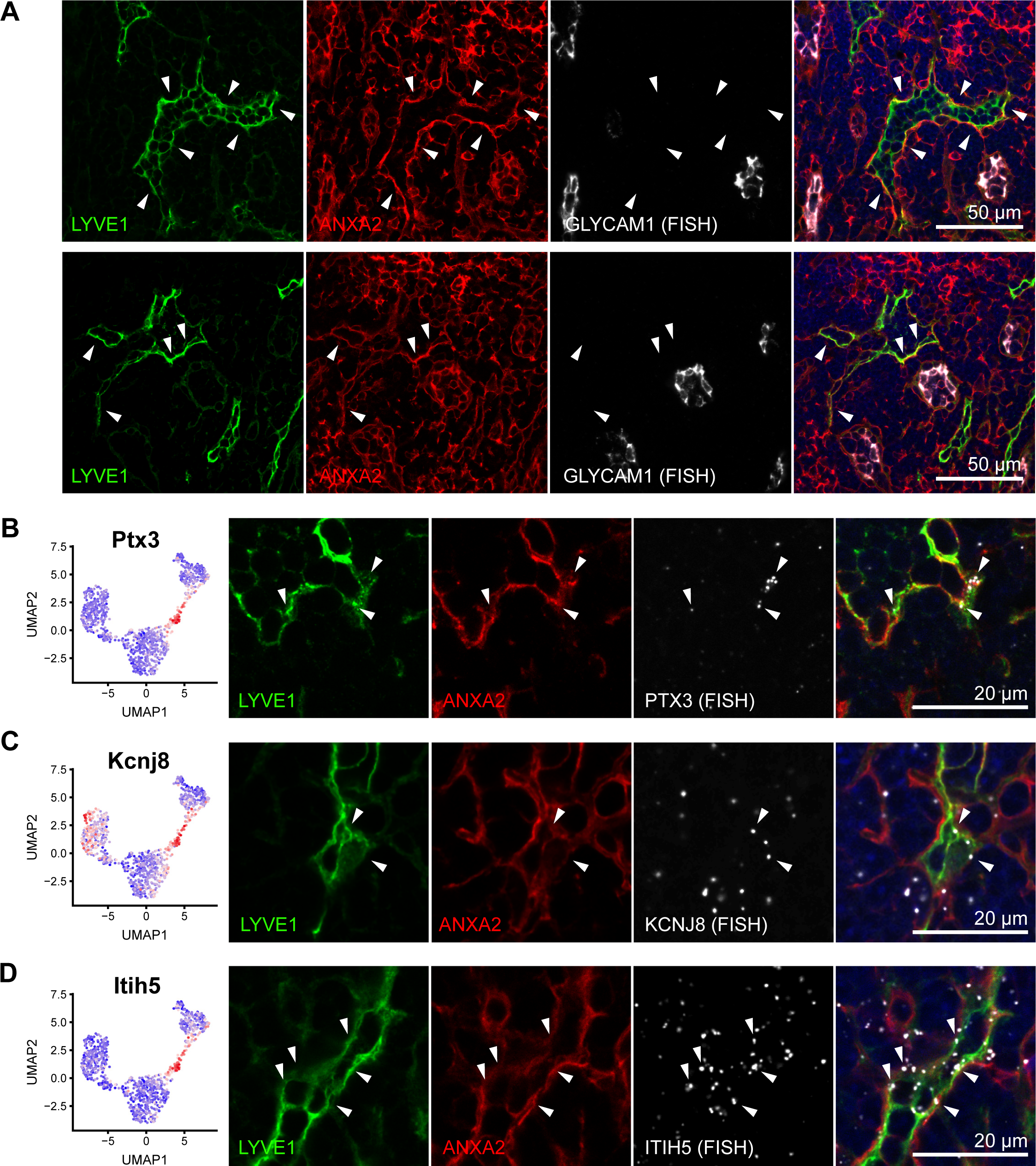
A unique subset of cortical and medullary sinuses. (A) Representative images of immunofluorescence staining for LYVE1 (green) and ANXA2 (red), in combination with RNA FISH to detect Glycam1 (white). ANXA2+ lymphatic sinuses are present in the cortex, close to the medulla (indicated by white arrowheads), and are often in close proximity to Glycam1+ HEVs. (B-D) Expression of the cluster 4 LEC marker genes Ptx3 (B), Kcnj8 (C) and Itih5 (D) by RNA sequencing (left panels) and RNA FISH (right panels). Immunofluorescence staining for ANXA2 (red) and LYVE1 (green) was used to highlight cluster 4 sinuses. White arrowheads point to LECs expressing Ptx3, Kcnj8 and Itih5 transcripts (white).

Previously, a subset of blind-ended sinuses has been described in the cortex of rat and mouse LNs that connect to medullary sinuses and may act as rapid egress structures for lymphocytes entering the LN through adjacent HEVs (Grigorova et al., 2010; He, 1985; Ohtani and Ohtani, 2008). Due to the close proximity of some cluster 4 sinuses to HEVs (Figure 7A), we hypothesized that they might be identical to those egress structures. To further investigate this hypothesis, we intravenously injected CFSE-labeled splenocytes into syngeneic recipient mice and analyzed their location in inguinal LNs after 10 and 30 min. In line with previously published data (Grigorova et al., 2010), we found that after 10 min, most infused immune cells were observed within HEVs (data not shown). 30 min after infusion however, many of the labeled cells had reached the LN parenchyma and eventually entered lymphatic sinuses. Using ANXA2 as a marker for cluster 4 sinuses as compared to medullary sinuses, we then quantified the percentage of infused splenocytes in each of the two sinus subtypes. Strikingly, infused cells were significantly more prevalent in ANXA2^+^ cortical sinuses than in ANXA2^−^ medullary sinuses (Figure 8A-B), although cortical sinuses were generally less frequent than medullary sinuses (data not shown). Together, these data further indicate that the cluster 4 LECs indeed represent the previously described lymphocyte egress structures in the LN cortex.

**Figure 8.**
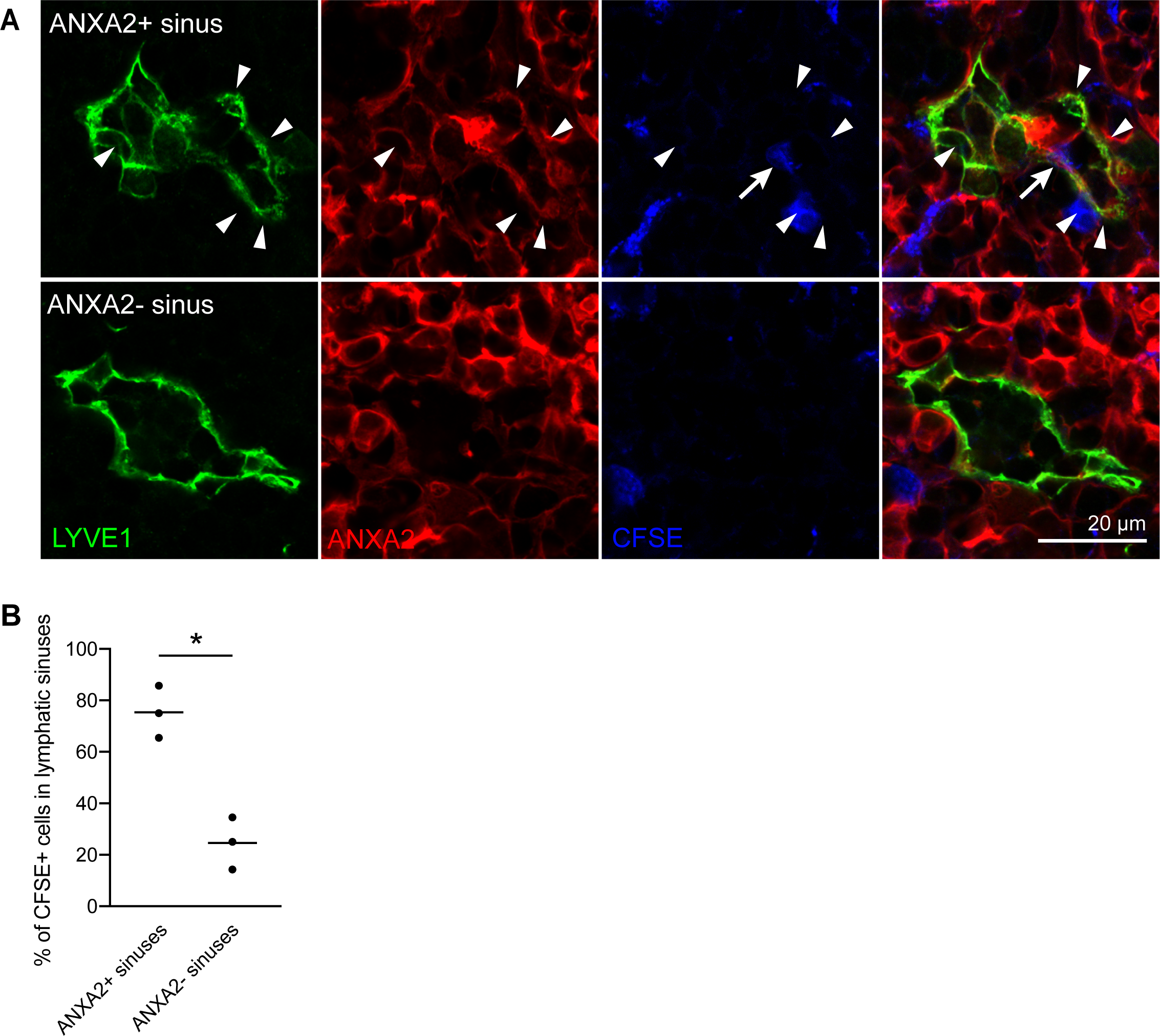
Lymphocytes egress from LNs via cluster 4 sinuses. CFSE-labelled splenocytes were infused via the tail vein and inguinal LNs were collected 30 min later. (A) Representative images of an ANXA2+ sinus (top) and an ANXA2-sinus (bottom), stained for LYVE1 (green) and ANXA2 (red). Arrowheads indicate ANXA2+ LECs. CFSE-labeled splenocytes (blue) eventually entered lymphatic sinuses (arrow). (B) Quantification of the percentage of CFSE+ splenocytes that entered lymphatic sinuses 30 min after infusion. CFSE-labeled cells were more frequently observed in ANXA2+ cluster 4 sinuses than in ANXA2-medullary sinuses. Each symbol represents one mouse (n = 3). * p < 0.05 (paired t-test).

## DISCUSSION

To unravel the phenotypic and functional heterogeneity of LN LECs, we performed single cell RNA sequencing coupled with unsupervised clustering, and identified at least four LEC subtypes that differed considerably in their transcriptome. Several previously described markers of LN LEC subsets residing in specific anatomical locations, such as ACKR4, LYVE1, and ITGA2B (Cordeiro et al., 2016; Ulvmar et al., 2014) were differentially expressed among these clusters, confirming the validity of our sequencing data and clustering approach. Most notably, the identification of new markers (Figure 9A) will enable the isolation of individual LN LEC subsets to study their transcriptomic alterations under pathological conditions in much greater details. The large number of differentially expressed genes furthermore implies that there are functional differences between LECs residing in different areas of the LN, which is most clearly seen in case of cLECs and fLECs lining the SCS. The fLECs function as a receptive surface for antigen-presenting cells entering the SCS with the afferent lymph (Braun et al., 2011; Ulvmar et al., 2014). In agreement with this, we found a significant enrichment of transcripts associated with inflammatory processes, including adhesion molecules such as CD44 and Glycam1, chemokines, and the innate-immunity related cochlin (Nystrom et al., 2018) in these cells (Supplementary Table 1). CLECs on the other hand expressed several matrix proteins which are likely involved in giving structural support to the LN and in providing a barrier towards the surrounding tissue. We also noted specific expression of PDGFs in cLECs, which probably mediate recruitment of perivascular supportive cells to the LN capsule (Bovay et al., 2018).

**Figure 9.**
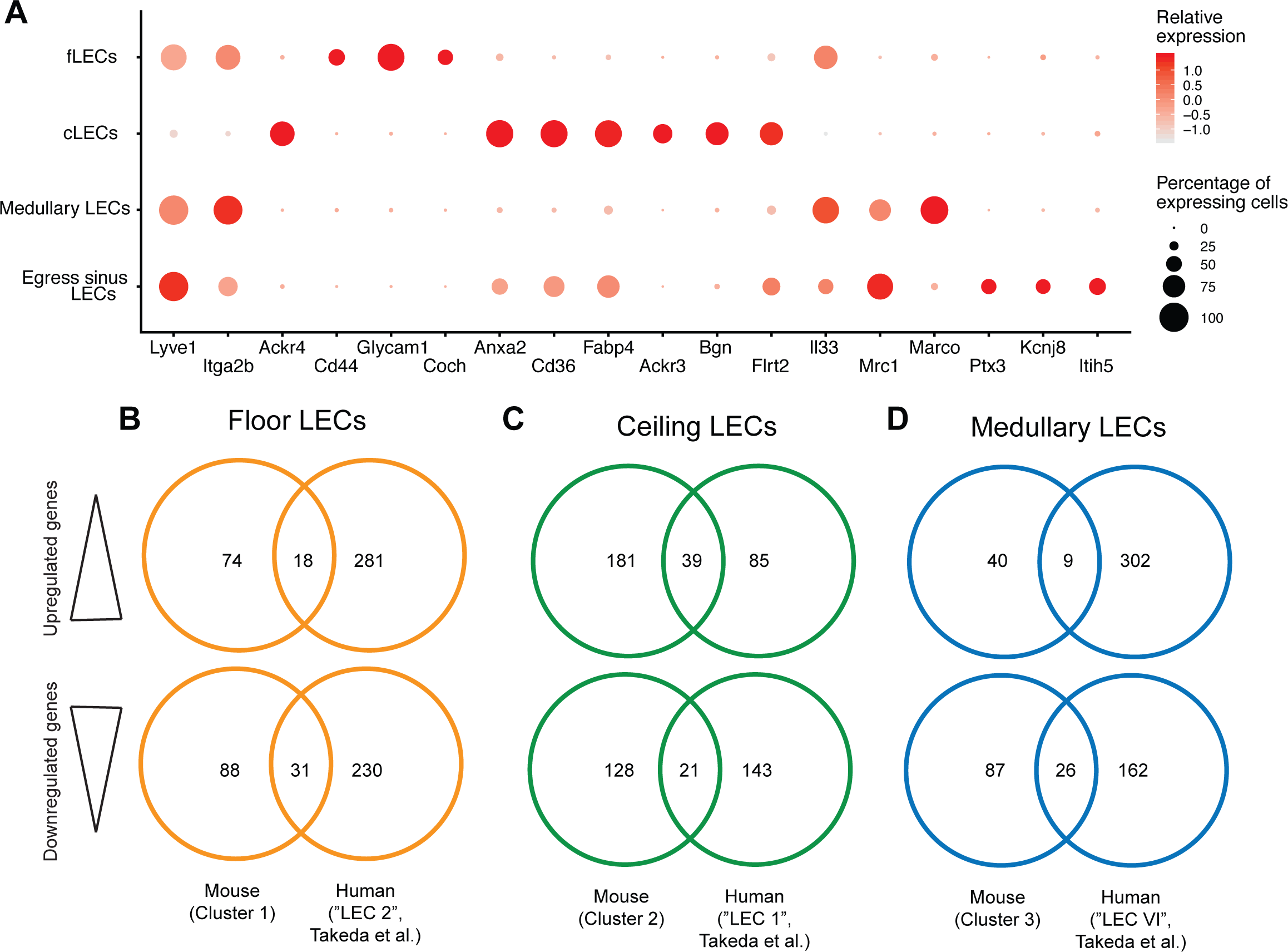
Overview of murine LN LEC subsets and comparison to human LN LECs. (A) Expression of the selected markers (x-axis) signifying individual LN LEC subsets (y-axis). Dot size denotes the proportion of cells with detectable expression. Color intensity indicates the relative mean expression level of the corresponding gene, using the original log-transformed counts. (B-D) Gene level comparison between mouse and human (Takeda et al., 2019) fLECs (B), cLECs (C), and medullary LECs (D), based on upregulated (top row) and downregulated (bottom) genes in each cluster compared to all other LECs. Venn diagrams display the number of differentially expressed genes that are shared or different between the two datasets.

Unexpectedly, we observed several proteins involved in the cellular uptake of modified LDL (Levitan et al., 2010) to be differentially expressed between cLECs and fLECs, namely CD36 (highly expressed in cLECs), MSR1 and FCGR2B (both excluded from cLECs) (Figure 3). While human lymph is basically devoid of very low-density lipoproteins, it does contain LDL (Reichl, 1990). In addition, the concentration of ApoB-protein is strongly reduced in efferent compared to afferent lymph in rats (Clement et al., 2018), suggesting that the lymphatic system may be involved in cholesterol transport and that LNs can actively remove LDL from the lymph. Our data using fluorescently-labeled, modified LDL suggest that LN LECs are at least partly responsible for the scavenging of lymphatic LDL. The difference between acetylated LDL, which was selectively taken up by cLECs, and oxidized LDL, which was selectively taken up by fLECs and cortical LECs, is probably due to differences in receptor affinities. It has been reported that MSR1 and FCGR2B require a high degree of LDL oxidation for efficient binding (Endemann et al., 1993), which is typically the case for commercially available oxidized LDL preparations, including the one used in the present study. Conversely, CD36 has a high affinity even for lowly oxidized LDL (Endemann et al., 1993), which may be reflected by the behavior of acetylated LDL in our assay. Clearly, further investigations are needed to examine the physiological significance of the capacity and selectivity of LDL uptake by LN LEC subtypes.

In 1985, Yechun He reported a subset of blind-ended lymphatic structures in proximity to HEVs in the inner cortex of rat mesenteric LNs, which he named “lymphatic labyrinth” (He, 1985). These structures were directly connected to medullary sinuses and supposedly act as an immediate egress portal for naïve lymphocytes entering the LN via the HEVs (Grigorova et al., 2010; He, 1985; Ohtani and Ohtani, 2008). Here, we have identified a transcriptionally distinct LN LEC subset that most likely represents the “lymphatic labyrinth” in mouse inguinal LNs. The LECs in this subset (cluster 4) shared high expression of several genes with either cLECs or medullary LECs, but also expressed several unique markers including PTX3, ITIH5, and KCNJ8, which we confirmed using *in situ* RNA hybridization. Lymphocyte egress from LNs depends on S1P gradients, which are established by LECs via expression of S1P kinases (SPHK1, SHPK2, (Pham et al., 2010)), S1P lyase (SGPL1, (Schwab et al., 2005)), and the sphingolipid transporter SPNS2 (Mendoza et al., 2012). In line with this, cluster 4 LECs expressed significantly more SPNS2 than other LN LECs, whereas SPHK1, SPHK2 and SGPL1 were expressed similarly in most LN LEC subsets (data not shown).

Previously, LN LECs have been implicated in the regulation of peripheral tolerance, expressing and presenting peripheral tissue self-antigens such as tyrosinase (TYR) (Cohen et al., 2010; Tewalt et al., 2012). The lack of co-stimulatory molecules and their constitutive expression of T cell-inhibitory molecules such as PD-L1, leads to the elimination of potential auto-reactive CD8^+^ T cells (Tewalt et al., 2012). Subsequently, PD-L1 expression has been mapped to subcapsular and medullary LECs by flow cytometry and immunofluorescence staining in mouse LNs, whereas TYR expression was predominant in medullary LECs (Cohen et al., 2014). In agreement with this, we found PD-L1 transcript expression in fLECs as well as medullary / cortical LECs, whereas TYR expression was highest in medullary LECs and LECs residing in the egress structures (cluster 4) (data not shown). This suggests that peripheral self-tolerance to TYR is primarily mediated by cortical and / or medullary LECs, while fLECs may be involved in tolerance towards other LEC-expressed self-antigens, or towards antigens taken up by fLECs from the lymph.

In a previous study, flow cytometry and laser-capture microdissection (LCM) were used to separate LECs in the SCS from other LN sinuses, followed by microarray analysis for transcriptional characterization (Iftakhar et al., 2016). However, it is unclear if LCM is sufficiently precise to separate individual sinus LECs from nearby cells such as sinus macrophages, and CD73 (NT5E), the marker used for FACS sorting of SCS versus other nodal LECs, was not differentially expressed in our single-cell RNA sequencing data (data not shown). Consequently, there was limited overlap between the transcriptional data presented in that study and our dataset (Iftakhar et al., 2016). More recently, Takeda et al. performed a single-cell RNA sequencing-based characterization of human LECs isolated from cervical and axillary LNs of tumor patients (Takeda et al., 2019). Similar to our study, the authors described 4 subsets of LN LECs within the nodes, including fLECs, cLECs, and medullary LECs. This suggests that these major LN LEC subtypes are conserved between species. A comparison between human (Takeda et al., 2019) and mouse fLECs, cLECs and medullary LECs based on up- and downregulated genes in those clusters (compared to all other LECs in the corresponding datasets) furthermore indicated transcriptional similarities, most prominently in cLECs (Figure 9B-D). This included expression of CD36, suggesting that human cLECs may also have the capacity to scavenge modified LDL. However, the majority of up- and downregulated genes were different in those LEC subsets, most likely due to the differences of the investigated tissues (species, anatomical location, disease status) and the technologies used (tissue digestion and LEC isolation, sequencing platforms, etc.). Furthermore, Takeda et al. failed to detect a cluster of cortical, lymphocyte egress-associated sinuses, possibly due to the 3’ transcriptomic profiling approach and the limited sequencing depth employed in their study (Takeda et al., 2019). On the other hand, we did not observe a specific medullary cLEC subpopulation as identified by Takeda et al., which therefore appears to be human specific.

Taken together, our data provide the first comprehensive transcriptional analysis of skin-draining LN LECs from naïve mice at the single cell level, identifying LEC subsets, new marker genes, and subset-specific functions. It will be of great interest to investigate, in future studies, the specific changes in LEC subset composition and gene expression patterns in pathological conditions such as inflammation and cancer. Moreover, whole transcriptome analysis of skin-draining LN LECs in comparison to those from e.g. cervical and mandibular nodes that drain the brain (Ma et al., 2017) or mesenteric nodes might provide new insights into the cellular basis of inter-nodal phenotypic and functional heterogeneity.

## MATERIAL AND METHODS

### Animals and ethics statement

C57Bl/6N and congenic Prox1-GFP reporter mice (Choi et al., 2011) were bred in house in an SOPF-level facility. Ackr4-GFP mice (Heinzel et al., 2007) were kindly provided by Prof. Cornelia Halin (Institute of Pharmaceutical Sciences, ETH Zurich). All in vivo experiments were approved by a local ethics committee (Kantonales Veterinäramt Zürich).

### Isolation of LN LECs

LECs were isolated from inguinal LNs of C57Bl/6N wildtype mice essentially as described before (Fletcher et al., 2011). In brief, the LNs were dissected and the capsule was broken using 23G injection needles. Subsequently, the tissue was digested in a solution containing 0.2 mg/ml Collagenase Type I (Worthington, Lakewood, NJ), 0.8 mg/ml Dispase II and 0.1 mg/ml DNAse I (both Roche, Basel, Switzerland) at 37°C. The samples were intermittently inverted or mixed by pipetting, and the entire digestion mix was renewed 2 times during the procedure. Once the tissue had completely dissolved, the cell suspension was washed with FACS buffer (1% FBS, 0.1 M EDTA, 0.02 % NaN_3_), blocked with anti-CD16/32 (101302, Biolegend, San Jose, CA), and labelled with anti-CD45.2-FITC (BD553772, BD Biosciences, San Diego, CA), anti-CD31-APC (BD551262, BD Biosciences) and anti-podoplanin-PE (12-5381-82, Thermo Fisher, Waltham, MA). Zombie-NIR (423106, Biolegend) was used for life/dead discrimination. Single, living CD45^−^ CD31^+^ podoplanin^+^ LECs were sorted directly into 384-well plates containing 0.8 µl of lysis buffer (0.1% Triton X-100, 2.5 mM dNTPs, 2.5 µM oligo-dT, 1 U/µl RNasin Plus RNase inhibitor (Promega, Madison, WI)) using a FACS Aria II instrument (BD Biosciences). Immediately after sorting, plates were centrifuged and stored at −80°C until further processing.

### Single-cell sequencing

Library preparation and sequencing were done at the Functional Genomic Center Zurich (FGCZ). In brief, the libraries were prepared using a miniaturised version of the Smart-seq2 protocol (Picelli et al., 2014) with the help of a Mosquito HV pipetting robot (TTP Labtech, Melbourn, UK). Reverse transcription was performed in a final volume of 2 µl followed by cDNA amplification in a final volume of 5 µl. The quality of the cDNAs was evaluated using a 2100 Bioanalyzer (Agilent, Santa Clara, CA). 0.1 ng of cDNA from each cell on the plate was individually tagmented using the Nextera XT kit (Illumina, San Diego, CA) in a final volume of 5 µl, followed by barcoding and library amplification in a final volume of 10 µl. The resulting 384 libraries were pooled, double-sided size selected (0.5x followed by 0.8x ratio using Ampure XP beads (Beckman Coulter, Brea, CA)) and quantified using a 4200 TapeStation System (Agilent). The pool of libraries was sequenced in Illumina HiSeq2500 using single-read 125 bp chemistry with a depth of around 750,000 reads per cell (around 300 Mio reads per plate).

### Data processing, unsupervised clustering and differential expression analyses

The Nextera adapter sequences and low quality bases were removed using trimmomatic v0.33 (Bolger et al., 2014). Trimmed reads were aligned to the Ensembl mm10 mouse reference genome (release 92) using STAR v2.4.2a (Dobin et al., 2013). Gene expression quantification was computed with the ‘featureCounts’ function in the Rsubread package v1.26.1 (Liao et al., 2019). Quality filtering was performed with the scran package v1.4.5, cells with library size or feature size 2.5 median absolute deviations (MADs) away from the median, or with mitochondrial contents 3 MADs above the median were dropped as outliers (Lun et al., 2016). Genes expressed in at least 15% of cells were grouped in accordance with their count-depth relationship by SCnorm v0.99.7, which applied a quantile regression within each group to estimate scaling factors and normalize for sequencing depth (Bacher et al., 2017). Cells with detected CD45 expression were removed before downstream analyses. To address the batch effects observed across different 384-well plates, we matched mutual nearest neighbors (MNNs) by applying the ‘fastMNN’ function implemented in the batchelor package v1.0.1 (Haghverdi et al., 2018). Gene-specific variance was decomposed into biological and technical components. Variable features were defined as the genes displaying positive biological variances (FDR < 0.05, mean expression level between 0.25 and 4). The variable features were then subjected to merged principle component analysis (PCA) across all batches. Unsupervised clustering was performed on the corrected low-dimensional coordinates (PC 1-10) using the Seurat package v2.3.4 (Satija et al., 2015) and visualized with Uniform Manifold Approximation and Projection (UMAP) (Becht et al., 2018). Differentially expressed genes in each cluster compared to all other clusters were identified by the ‘FindMarkers’ function (min.pct = 0.20, logfc.threshold = 0.6, p_val_adj < 0.01) using the MAST test (Finak et al., 2015). The expression patterns of selected markers were plotted by the ‘FeaturePlot’ function using the corrected expression values. The entire gene expression data will be made accessible via a public repository.

### Analysis and comparison of previously published human LN LEC data

Raw data were downloaded from GSE124494 and re-analyzed with Seurat. Quality control was performed as previously described (Takeda et al., 2019). Six human LN datasets were aligned using Canonical Correlation Analysis (CCA) with highly variable genes identified in at least 2 datasets. Clusters were defined using the aligned canonical correlation vectors (CC) 1-30 and resolution 0.5. Only the LEC population was subsetted for downstream analysis. Differentially expressed genes among clusters were identified with ‘FindAllMarkers (min.pct = 0.25, logfc.threshold = 0.25 and p_val_adj < 0.05). Genes upregulated in LEC I, II and VI were compared to genes upregulated in our cLEC, fLEC and medullary LEC clusters, respectively. Orthologous genes were converted using the biomaRt package (Durinck et al., 2009).

### Immunofluorescence staining

Inguinal LNs of C57Bl/6N wildtype mice were dissected, embedded in OCT compound and frozen in liquid nitrogen. In the case of Ackr4-GFP reporter mice, inguinal, mandibular, iliac and mesenteric LNs were dissected, fixed with 2% paraformaldehyde for 2 h at room temperature, treated with 1 M sucrose overnight, embedded in OCT compound and frozen in liquid nitrogen. 7 µm sections were cut with a cryostat, fixed in ice-cold acetone and 80% methanol, rehydrated in PBS, and subsequently blocked in PBS + 0.2% BSA, 5% donkey serum, 0.3% Triton-X100, and 0.05% NaN_3_ (blocking solution). Primary antibodies included goat anti-LYVE1 (AF2125, R&D, Minneapolis, MN), rabbit anti-LYVE1 (11-034, AngioBio, San Diego, CA), rat anti-LYVE1 (NBP1-43411, Novus Biologicals, Centennial, CO), goat anti-prox1 (AF2727, R&D), rabbit anti-CD3 (NB600-1441, Novus Biologicals), rat anti-IgD (1120-01, SouthernBiotech, Birmingham), rat anti-CD4 (BD553647, BD Biosciences), rat anti-F4/80 (MCA487R, Bio-Rad, Hercules, CA), rat anti-CD169 (MCA884, Bio-Rad), rat anti-ITGA2B (BD553847, BD Bioscience), rat anti-CD44 (103002, Biolegend), rabbit anti-ANXA2 (ab178677, Abcam, Cambridge, UK), rabbit anti-FABP4 (15872-1-AP, ProteinTech, Rosemont, IL), goat anti-CD36 (AF2519, R&D), goat anti-IL33 (AF3626, R&D), goat anti-MRC1 (AF2535, R&D), rat anti-MARCO (GTX39744, Genetex, Irvine, CA), rat anti-MADCAM1 (BD553805, BD Bioscience), rabbit anti-BGN (HPA003157, Sigma, St. Louis, MO), and goat anti-FLRT2 (AF2877, R&D). They were suspended in blocking solution and incubated at 4°C overnight, followed by incubation with Alexa488, Alexa594, or Alexa647-conjugated secondary antibodies (donkey anti-rat, donkey anti-rabbit, donkey anti-goat, all Thermo Fisher) together with Hoechst33342 (Sigma) for nuclear counterstaining. Images were captured with an LSM 780 upright confocal microscope (Zeiss, Jena, Germany) and analyzed with Fiji (Schindelin et al., 2012).

### RNA-FISH

RNA fluorescence *in situ* hybridizations (FISH) were performed using the RNAscope Multiplex Fluorescent Reagent Kit v2 (ACD, Newark, CA) according to the manufacturer’s instructions. Inguinal LNs were fixed with 10% neutral-buffered paraformaldehyde and embedded in paraffin for sectioning (7 µm). Antigen retrieval was performed with Target Retrieval Reagent for 15 min after deparaffinization. Slides were treated with Protease Plus for 30 min, incubated with probes in the ACD HybEZ hybridization system and stained with Opal 570. The following RNAscope probes were used: Ackr3 (C1), Btln9 (C1), Coch (C3), Glycam1 (C1), Itih5 (C1), Kcnj8 (C1), and Ptx3 (C1). Slides were stained with primary antibodies as described above, followed by incubation with Alexa488 and/or Alexa647-conjugated secondary antibodies together with Hoechst33342 for nuclear counterstaining, and mounted with ProLong Gold Antifade Mounting medium (Thermo Fisher). Images were captured with an LSM 780 upright confocal microscope and analyzed with Fiji.

### Flow cytometry analysis

Inguinal LNs from C57Bl/6N wildtype and Ackr4-GFP where processed, washed and Fc-blocked as described above (“Isolation of LN LECs”). Subsequently, cell suspensions were stained with anti-CD45-PacificBlue (103126, Biolegend), anti-podoplanin-PE, anti-CD31-PerCp/Cy5.5 (102522, Biolegend), anti-ITGA2B-BV421 (133911, Biolegend), anti-MRC1-APC (141708, Biolegend), anti CD44-BV650 (103049, Biolegend), and Zombie-NIR, and analyzed on a LSRFortessa flow cytometer (BD Biosciences).

### Light sheet microscopy

Inguinal LNs derived from Prox1-GFP reporter mice were fixed with 4% paraformaldehyde for 2 h at room temperature, permeabilized in 0.5% Triton X-100 in PBS for 2 days, and stained with chicken anti-GFP (GFP1010, Aves Labs, Davis, CA) and rabbit anti-ANXA2 for 7 days followed by incubation with Alexa488 and Alexa594-conjugated secondary antibodies for 7 days. Optical clearing was performed as described previously (Commerford et al., 2018). In brief, stained LNs were embedded in 1% ultrapure LMP Agarose (Thermo Fisher) on ice, dehydrated in a series of 50%, 70%, 95%, and 100% methanol, pre-cleared with 50% BABB (benzyl alcohol/ benzyl benzoate 1:2) in methanol followed by clearing in 100% BABB overnight. Wholemount images were captured with an UltraMicroscope I (LaVision Biotec, Bielefeld, Germany) and analyzed with Fiji.

### In vivo LDL tracing assay

10 µg of Dil-labeled human acetylated LDL (Kalen Biomedical, Germantown, MD) or 10 µg of Dil-labeled human oxidized LDL (Thermo Fisher) was intradermally injected unilaterally close to the base of the tail of Ackr4-GFP reporter mice under isoflurane anesthesia. An equal volume of PBS was injected on the opposite side as control. Draining inguinal LNs were collected 1 h later, fixed with 2% paraformaldehyde for 2 h at room temperature, embedded in OCT compound and frozen in liquid nitrogen. Immunofluorescence staining using a goat anti-LYVE1 antibody (R&D) followed by incubation with an Alexa647-conjugated secondary antibody together with Hoechst33342 (Sigma) was performed on 7 µm sections without acetone fixation. Images covering the whole SCS and SCS/CS regions were captured with an LSM 780 upright confocal microscope. The intensity of LDL staining in the ACKR4^+^ area and the ACKR4^−^ LYVE1^+^ area was measured with Fiji. For quantification, the average signal intensity of all images representing each individual mouse was normalized (ACKR4^−^ LYVE1^+^ area = 1).

### Adoptive lymphocyte transfer and tracing

Splenocytes were collected from naïve C57Bl/6N wildtype mice after lysis of red blood cells with PharmLyse buffer (BD Bioscience) and were labeled with 5 mM of carboxy-fluorescein diacetate succinimidyl ester (CFSE) (Sigma) in PBS for 15 min at 37 °C. 2×10^6^ labeled splenocytes were infused into the tail vein of sex-matched recipient mice. Inguinal LNs were collected 30 min later, fixed with 10% paraformaldehyde overnight at room temperature, and embedded in paraffin. 7 µm and 40 µm sections were deparaffinized, followed by antigen retrieval (10 mM citrate buffer, pH 6.0) and immunofluorescence staining using goat anti-LYVE1 (R&D) and rabbit anti-ANXA2 (Abcam) antibodies and donkey anti-goat and anti-rabbit Alexa594 and Alexa647-conjugated secondary antibodies. Images covering all cortical and medullary regions were captured with LSM 780 upright confocal microscope and analyzed with Fiji. The number of CFSE-labeled cells in ANXA2^+^ sinuses and in ANXA2^−^sinuses were counted manually using 7 µm sections. Maximum intensity projections of confocal z-stacks images were prepared using 40 µm sections of the same samples.

## Statistical analysis

Statistical analysis was performed using GraphPad Prism (GraphPad Software, San Diego, CA). Student’s *t*-test was used for comparisons of two groups. A *p*-value < 0.05 was considered statistically significant. ScRNA-seq data analyses and graphical interpretation were performed using R v3.6.1.

## Supplementary data

Supplementary data comprise Supplementary Table 1 (gene ontology for 4 LEC subtypes) and Supplementary Figures 1-5 with additional immunofluorescence staining and FACS data.

## AUTHOR CONTRIBUTIONS

NF designed and performed experiments, analyzed data, and wrote the manuscript; YH analyzed and visualized RNA-seq data, and wrote the manuscript; MDA and CT designed and performed experiments; MD conceptualized the study, designed experiments, and revised the manuscript. LCD conceptualized the study, designed and performed experiments, analyzed data, and wrote the manuscript.

## ACKNOWLEDGEMENTS

The authors gratefully acknowledge excellent technical and experimental support by Jeannette Scholl (ETH Zurich), Nikola Cousin (ETH Zurich), Dr. Steven Proulx (University of Berne) and Dr. Emilio Yanguez (FGCZ Zurich). Ackr4-GFP mice were bred and kindly provided by Mona Friess and Prof. Cornelia Halin (ETH Zurich). This work was supported by grants from Krebsliga Zurich and the Vontobel Foundation (to LCD) and by Swiss National Science Foundation grants 310030_166490 and 310030B_185392, and European Research Council grant LYVICAM (to MD).

## SUPPLEMENTARY MATERIAL

**Supplementary Table 1**

Gene ontology analysis of biological process (GO_BP) terms using differentially expressed genes among the 4 LN LEC clusters. Only the top 10 most significantly enriched terms are shown for each of the clusters.

**Supplementary Figure 1**

(A) Immunofluorescence staining for LYVE1 (green) and MADCAM1 (red), showing specific MADCAM1 staining in the floor of the subcapsular sinus. (B) Immunofluorescence staining for LYVE1 (green) and ITGA2B (red). LYVE1 and ITGA2B were clearly detectable in fLECs and cortical LECS both in the SCS and the IF-SCS regions. (C, D) Immunofluorescence staining for LYVE1 (green) in Ackr4-GFP reporter mice. ACKR4+ cLECs (white) were detected both in the SCS region and the IF-SCS region.

**Supplementary Figure 2**

(A) Light sheet fluorescence microscopy image of an optically cleared inguinal LN derived from a Prox1-GFP reporter mouse. Immunofluorescence staining for ANXA2 (red) revealed that afferent lymphatic collectors express ANXA2 (white arrowhead). (B, C) Expression of new cLEC / cluster 2 marker genes BGN (B) and FLRT2 (C) by RNA sequencing (left panels) and immunofluorescence staining (right panels) in Ackr4-GFP reporter mice. GFP (white) and immunofluorescence co-staining for LYVE1 (green) served as markers for cLECs and fLECs, respectively.

**Supplementary Figure 3**

(A-D) Immunofluorescence images of mandibular, iliac and mesenteric LN sections derived from Ackr4-GFP reporter mice, stained for LYVE1 (green), CD44 (red) (A), ANXA2 (red) (B), CD36 (red) (C) or MRC1 (red) (D). GFP fluorescence is shown in white.

**Supplementary Figure 4**

(A) Gating strategy to identify the major LEC subsets (cLECs, fLECs, medullary LECs) in inguinal LNs from Ackr4-GFP mice by flow cytometry. Within LN stromal cells (pregated as CD45-, Zombie-NIR-singlets), LECs were identified as podoplanin (PDPN)+ CD31+ cells. cLECs were identified by GFP expression. Among the remaining cells, medullary LECs expressed MRC1 and were predominantly ITGA2B+, whereas fLECs were MRC1-but expressed relatively high levels of CD44 and ITGA2B. (B) Staining controls for GFP (using wildtype C57Bl/6N mice), MRC1, ITGA2B and CD44. (C) Intensity histograms for GFP, MRC1, ITGA2B and CD44 in cLECs (green curve), fLECs (orange curve) and medullary LECs (blue curve) identified as shown in panel A.

**Supplementary Figure 5**

(A-D) Representative images of cluster 4 sinuses stained for LYVE1 (green) and ANXA2 (red). The location relative to major immune cell populations is shown by staining for IgD (A), CD4 (B), F4/80 (C) and CD169 (D). (D-G) Immunofluorescence staining for LYVE1 (green), ANXA2 (red), and PROX1 (A), MRC1 (B) and MARCO (C) (white). White arrowheads indicate LYVE1+ / ANXA2+ cells.

## REFERENCES

Abeler-Dorner, L., M. Swamy, G. Williams, A.C. Hayday, and A. Bas. 2012. Butyrophilins: an emerging family of immune regulators. Trends Immunol. 33:34–41.

Angeli, V., F. Ginhoux, J. Llodra, L. Quemeneur, P.S. Frenette, M. Skobe, R. Jessberger, M. Merad, and G.J. Randolph. 2006. B cell-driven lymphangiogenesis in inflamed lymph nodes enhances dendritic cell mobilization. Immunity 24:203–215.

Bacher, R., L.F. Chu, N. Leng, A.P. Gasch, J.A. Thomson, R.M. Stewart, M. Newton, and C. Kendziorski. 2017. SCnorm: robust normalization of single-cell RNA-seq data. Nat. Methods 14:584–586.

Banerji, S., J. Ni, S.X. Wang, S. Clasper, J. Su, R. Tammi, M. Jones, and D.G. Jackson. 1999. LYVE-1, a new homologue of the CD44 glycoprotein, is a lymph-specific receptor for hyaluronan. J. Cell Biol. 144:789–801.

Becht, E., L. McInnes, J. Healy, C.A. Dutertre, I.W.H. Kwok, L.G. Ng, F. Ginhoux, and E.W. Newell. 2019. Dimensionality reduction for visualizing single-cell data using UMAP. Nat. Biotechnol. 37:38–44.

Bolger, A.M., M. Lohse, and B. Usadel. 2014. Trimmomatic: a flexible trimmer for Illumina sequence data. Bioinformatics 30:2114–2120.

Bovay, E., A. Sabine, B. Prat-Luri, S. Kim, K. Son, A.H. Willrodt, C. Olsson, C. Halin, F. Kiefer, C. Betsholtz, N.L. Jeon, S.A. Luther, and T.V. Petrova. 2018. Multiple roles of lymphatic vessels in peripheral lymph node development. J. Exp. Med. 215:2760–2777.

Braun, A., T. Worbs, G.L. Moschovakis, S. Halle, K. Hoffmann, J. Bolter, A. Munk, and R. Forster. 2011. Afferent lymph-derived T cells and DCs use different chemokine receptor CCR7-dependent routes for entry into the lymph node and intranodal migration. Nat. Immunol. 12:879–887.

Choi, I., H.K. Chung, S. Ramu, H.N. Lee, K.E. Kim, S. Lee, J. Yoo, D. Choi, Y.S. Lee, B. Aguilar, and Y.K. Hong. 2011. Visualization of lymphatic vessels by Prox1-promoter directed GFP reporter in a bacterial artificial chromosome-based transgenic mouse. Blood 117:362–365.

Clement, C.C., W. Wang, M. Dzieciatkowska, M. Cortese, K.C. Hansen, A. Becerra, S. Thangaswamy, I. Nizamutdinova, J.Y. Moon, L.J. Stern, A.A. Gashev, D. Zawieja, and L. Santambrogio. 2018. Quantitative profiling of the lymph node clearance capacity. Sci. Rep. 8:11253.

Cohen, J.N., C.J. Guidi, E.F. Tewalt, H. Qiao, S.J. Rouhani, A. Ruddell, A.G. Farr, K.S. Tung, and V.H. Engelhard. 2010. Lymph node-resident lymphatic endothelial cells mediate peripheral tolerance via Aire-independent direct antigen presentation. J. Exp. Med. 207:681–688.

Cohen, J.N., E.F. Tewalt, S.J. Rouhani, E.L. Buonomo, A.N. Bruce, X. Xu, S. Bekiranov, Y.X. Fu, and V.H. Engelhard. 2014. Tolerogenic properties of lymphatic endothelial cells are controlled by the lymph node microenvironment. PLoS One 9:e87740.

Commerford, C.D., L.C. Dieterich, Y. He, T. Hell, J.A. Montoya-Zegarra, S.F. Noerrelykke, E. Russo, M. Rocken, and M. Detmar. 2018. Mechanisms of tumor-induced lymphovascular niche formation in draining lymph nodes. Cell Rep. 25:3554–3563 e3554.

Cordeiro, O.G., M. Chypre, N. Brouard, S. Rauber, F. Alloush, M. Romera-Hernandez, C. Benezech, Z. Li, A. Eckly, M.C. Coles, A. Rot, H. Yagita, C. Leon, B. Ludewig, T. Cupedo, F. Lanza, and C.G. Mueller. 2016. Integrin-alpha IIb identifies murine lymph node lymphatic endothelial cells responsive to RANKL. PLoS One 11:e0151848.

Cyster, J.G., and S.R. Schwab. 2012. Sphingosine-1-phosphate and lymphocyte egress from lymphoid organs. Annu. Rev. Immunol. 30:69–94.

Dieterich, L.C., and M. Detmar. 2016. Tumor lymphangiogenesis and new drug development. Adv. Drug Deliv. Rev. 99:148–160.

Dieterich, L.C., C.D. Seidel, and M. Detmar. 2014. Lymphatic vessels: new targets for the treatment of inflammatory diseases. Angiogenesis 17:359–371.

Dobin, A., C.A. Davis, F. Schlesinger, J. Drenkow, C. Zaleski, S. Jha, P. Batut, M. Chaisson, and T.R. Gingeras. 2013. STAR: ultrafast universal RNA-seq aligner. Bioinformatics 29:15–21.

Durinck, S., P.T. Spellman, E. Birney, and W. Huber. 2009. Mapping identifiers for the integration of genomic datasets with the R/Bioconductor package biomaRt. Nat. Protoc. 4:1184–1191.

Endemann, G., L.W. Stanton, K.S. Madden, C.M. Bryant, R.T. White, and A.A. Protter. 1993. CD36 is a receptor for oxidized low density lipoprotein. J. Biol. Chem. 268:11811–11816.

Finak, G., A. McDavid, M. Yajima, J. Deng, V. Gersuk, A.K. Shalek, C.K. Slichter, H.W. Miller, M.J. McElrath, M. Prlic, P.S. Linsley, and R. Gottardo. 2015. MAST: a flexible statistical framework for assessing transcriptional changes and characterizing heterogeneity in single-cell RNA sequencing data. Genome Biol. 16:278.

Fletcher, A.L., D. Malhotra, S.E. Acton, V. Lukacs-Kornek, A. Bellemare-Pelletier, M. Curry, M. Armant, and S.J. Turley. 2011. Reproducible isolation of lymph node stromal cells reveals site-dependent differences in fibroblastic reticular cells. Front. Immunol. 2:35.

Gregory, J.L., A. Walter, Y.O. Alexandre, J.L. Hor, R. Liu, J.Z. Ma, S. Devi, N. Tokuda, Y. Owada, L.K. Mackay, G.K. Smyth, W.R. Heath, and S.N. Mueller. 2017. Infection programs sustained lymphoid stromal cell responses and shapes lymph node remodeling upon secondary challenge. Cell Rep. 18:406–418.

Grigorova, I.L., M. Panteleev, and J.G. Cyster. 2010. Lymph node cortical sinus organization and relationship to lymphocyte egress dynamics and antigen exposure. Proc. Natl. Acad. Sci. U. S. A. 107:20447–20452.

Haghverdi, L., A.T.L. Lun, M.D. Morgan, and J.C. Marioni. 2018. Batch effects in single-cell RNA-sequencing data are corrected by matching mutual nearest neighbors. Nat. Biotechnol. 36:421–427.

He, Y. 1985. Scanning electron microscope studies of the rat mesenteric lymph node with special reference to high-endothelial venules and hitherto unknown lymphatic labyrinth. Arch. Histol. Jpn. 48:1–15.

Heinzel, K., C. Benz, and C.C. Bleul. 2007. A silent chemokine receptor regulates steady-state leukocyte homing in vivo. Proc. Natl. Acad. Sci. U. S. A. 104:8421–8426.

Hirakawa, S., L.F. Brown, S. Kodama, K. Paavonen, K. Alitalo, and M. Detmar. 2007. VEGF-C-induced lymphangiogenesis in sentinel lymph nodes promotes tumor metastasis to distant sites. Blood 109:1010–1017.

Hirosue, S., E. Vokali, V.R. Raghavan, M. Rincon-Restrepo, A.W. Lund, P. Corthesy-Henrioud, F. Capotosti, C. Halin Winter, S. Hugues, and M.A. Swartz. 2014. Steady-state antigen scavenging, cross-presentation, and CD8+ T cell priming: a new role for lymphatic endothelial cells. J. Immunol. 192:5002–5011.

Iftakhar, E.K.I., R. Fair-Makela, A. Kukkonen-Macchi, K. Elima, M. Karikoski, P. Rantakari, M. Miyasaka, M. Salmi, and S. Jalkanen. 2016. Gene-expression profiling of different arms of lymphatic vasculature identifies candidates for manipulation of cell traffic. Proc. Natl. Acad. Sci. U. S. A. 113:10643–10648.

Imai, Y., M.S. Singer, C. Fennie, L.A. Lasky, and S.D. Rosen. 1991. Identification of a carbohydrate-based endothelial ligand for a lymphocyte homing receptor. J. Cell Biol. 113:1213–1221.

Kahari, L., R. Fair-Makela, K. Auvinen, P. Rantakari, S. Jalkanen, J. Ivaska, and M. Salmi. 2019. Transcytosis route mediates rapid delivery of intact antibodies to draining lymph nodes. J. Clin. Invest. 129:3086–3102.

Karaman, S., and M. Detmar. 2014. Mechanisms of lymphatic metastasis. J. Clin. Invest. 124:922–928.

Levitan, I., S. Volkov, and P.V. Subbaiah. 2010. Oxidized LDL: diversity, patterns of recognition, and pathophysiology. Antioxid. Redox. Signal. 13:39–75.

Liao, Y., G.K. Smyth, and W. Shi. 2019. The R package Rsubread is easier, faster, cheaper and better for alignment and quantification of RNA sequencing reads. Nucleic Acids Res. 47:e47.

Lun, A.T., D.J. McCarthy, and J.C. Marioni. 2016. A step-by-step workflow for low-level analysis of single-cell RNA-seq data with Bioconductor. F1000Res. 5:2122.

Ma, Q., B.V. Ineichen, M. Detmar, and S.T. Proulx. 2017. Outflow of cerebrospinal fluid is predominantly through lymphatic vessels and is reduced in aged mice. Nat. Commun. 8:1434.

Malhotra, D., A.L. Fletcher, J. Astarita, V. Lukacs-Kornek, P. Tayalia, S.F. Gonzalez, K.G. Elpek, S.K. Chang, K. Knoblich, M.E. Hemler, M.B. Brenner, M.C. Carroll, D.J. Mooney, S.J. Turley, and C. Immunological Genome Project. 2012. Transcriptional profiling of stroma from inflamed and resting lymph nodes defines immunological hallmarks. Nat. Immunol. 13:499–510.

Martens, J.H., J. Kzhyshkowska, M. Falkowski-Hansen, K. Schledzewski, A. Gratchev, U. Mansmann, C. Schmuttermaier, E. Dippel, W. Koenen, F. Riedel, M. Sankala, K. Tryggvason, L. Kobzik, G. Moldenhauer, B. Arnold, and S. Goerdt. 2006. Differential expression of a gene signature for scavenger/lectin receptors by endothelial cells and macrophages in human lymph node sinuses, the primary sites of regional metastasis. J. Pathol. 208:574–589.

Mendoza, A., B. Breart, W.D. Ramos-Perez, L.A. Pitt, M. Gobert, M. Sunkara, J.J. Lafaille, A.J. Morris, and S.R. Schwab. 2012. The transporter Spns2 is required for secretion of lymph but not plasma sphingosine-1-phosphate. Cell Rep. 2:1104–1110.

Nystrom, A., O. Bornert, T. Kuhl, C. Gretzmeier, K. Thriene, J. Dengjel, A. Pfister-Wartha, D. Kiritsi, and L. Bruckner-Tuderman. 2018. Impaired lymphoid extracellular matrix impedes antibacterial immunity in epidermolysis bullosa. Proc. Natl. Acad. Sci. U. S. A. 115:E705–E714.

Ohtani, O., and Y. Ohtani. 2008. Structure and function of rat lymph nodes. Arch. Histol. Cytol. 71:69–76.

Pham, T.H., P. Baluk, Y. Xu, I. Grigorova, A.J. Bankovich, R. Pappu, S.R. Coughlin, D.M. McDonald, S.R. Schwab, and J.G. Cyster. 2010. Lymphatic endothelial cell sphingosine kinase activity is required for lymphocyte egress and lymphatic patterning. J. Exp. Med. 207:17–27.

Picelli, S., O.R. Faridani, A.K. Bjorklund, G. Winberg, S. Sagasser, and R. Sandberg. 2014. Full-length RNA-seq from single cells using Smart-seq2. Nat. Protoc. 9:171–181.

Rantakari, P., K. Auvinen, N. Jappinen, M. Kapraali, J. Valtonen, M. Karikoski, H. Gerke, E.K.I. Iftakhar, J. Keuschnigg, E. Umemoto, K. Tohya, M. Miyasaka, K. Elima, S. Jalkanen, and M. Salmi. 2015. The endothelial protein PLVAP in lymphatics controls the entry of lymphocytes and antigens into lymph nodes. Nat. Immunol. 16:386–396.

Reichl, D. 1990. Lipoproteins of human peripheral lymph. Eur. Heart J. 11 Suppl E:230–236.

Reynoso, G.V., A.S. Weisberg, J.P. Shannon, D.T. McManus, L. Shores, J.L. Americo, R.V. Stan, J.W. Yewdell, and H.D. Hickman. 2019. Lymph node conduits transport virions for rapid T cell activation. Nat. Immunol. 20:602–612.

Rodda, L.B., E. Lu, M.L. Bennett, C.L. Sokol, X. Wang, S.A. Luther, B.A. Barres, A.D. Luster, C.J. Ye, and J.G. Cyster. 2018. Single-Cell RNA Sequencing of lymph node stromal cells reveals niche-associated heterogeneity. Immunity 48:1014–1028 e1016.

Satija, R., J.A. Farrell, D. Gennert, A.F. Schier, and A. Regev. 2015. Spatial reconstruction of single-cell gene expression data. Nat. Biotechnol. 33:495–502.

Schindelin, J., I. Arganda-Carreras, E. Frise, V. Kaynig, M. Longair, T. Pietzsch, S. Preibisch, C. Rueden, S. Saalfeld, B. Schmid, J.Y. Tinevez, D.J. White, V. Hartenstein, K. Eliceiri, P. Tomancak, and A. Cardona. 2012. Fiji: an open-source platform for biological-image analysis. Nat. Methods 9:676–682.

Schwab, S.R., J.P. Pereira, M. Matloubian, Y. Xu, Y. Huang, and J.G. Cyster. 2005. Lymphocyte sequestration through S1P lyase inhibition and disruption of S1P gradients. Science 309:1735–1739.

Takeda, A., M. Hollmen, D. Dermadi, J. Pan, K.F. Brulois, R. Kaukonen, T. Lonnberg, P. Bostrom, I. Koskivuo, H. Irjala, M. Miyasaka, M. Salmi, E.C. Butcher, and S. Jalkanen. 2019. Single-cell survey of human lymphatics unveils marked endothelial cell heterogeneity and mechanisms of homing for neutrophils. Immunity 51:561–572 e565.

Tewalt, E.F., J.N. Cohen, S.J. Rouhani, C.J. Guidi, H. Qiao, S.P. Fahl, M.R. Conaway, T.P. Bender, K.S. Tung, A.T. Vella, A.J. Adler, L. Chen, and V.H. Engelhard. 2012. Lymphatic endothelial cells induce tolerance via PD-L1 and lack of costimulation leading to high-level PD-1 expression on CD8 T cells. Blood 120:4772–4782.

Ulvmar, M.H., K. Werth, A. Braun, P. Kelay, E. Hub, K. Eller, L. Chan, B. Lucas, I. Novitzky-Basso, K. Nakamura, T. Rulicke, R.J. Nibbs, T. Worbs, R. Forster, and A. Rot. 2014. The atypical chemokine receptor CCRL1 shapes functional CCL21 gradients in lymph nodes. Nat. Immunol. 15:623–630.

