## Supplementary figures and images for "Single-cell mapping reveals new markers and functions of lymphatic endothelial cells in lymph nodes"

### Supplementary Figures S1-S5

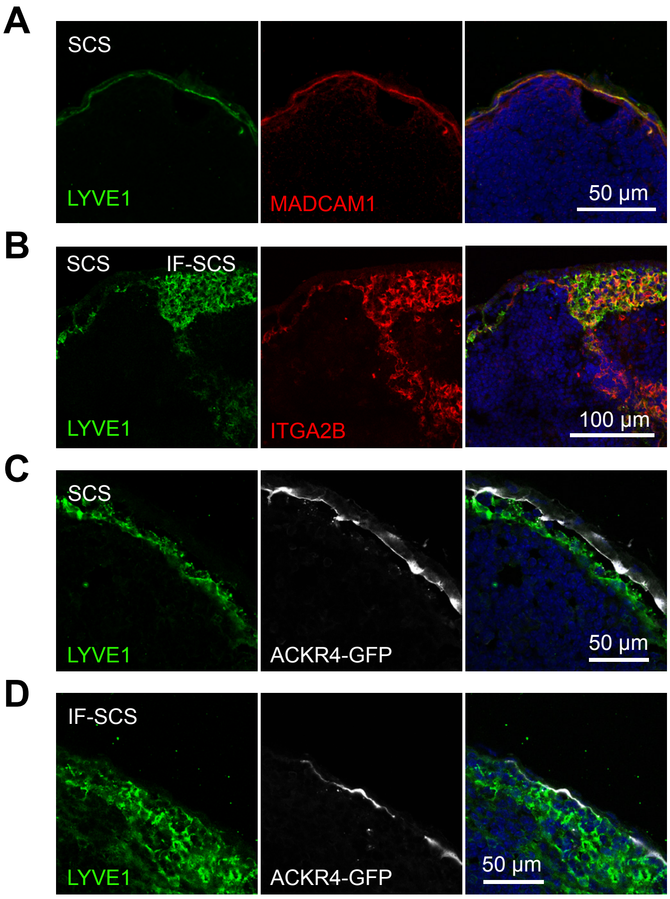

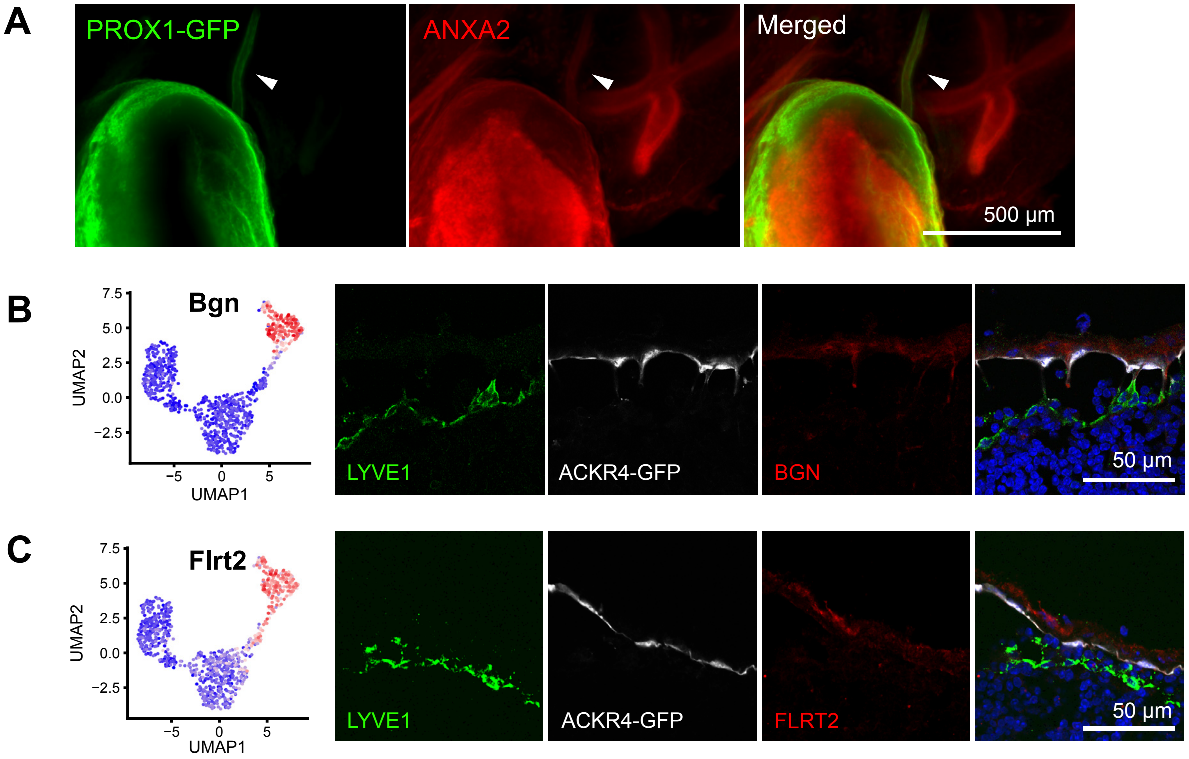

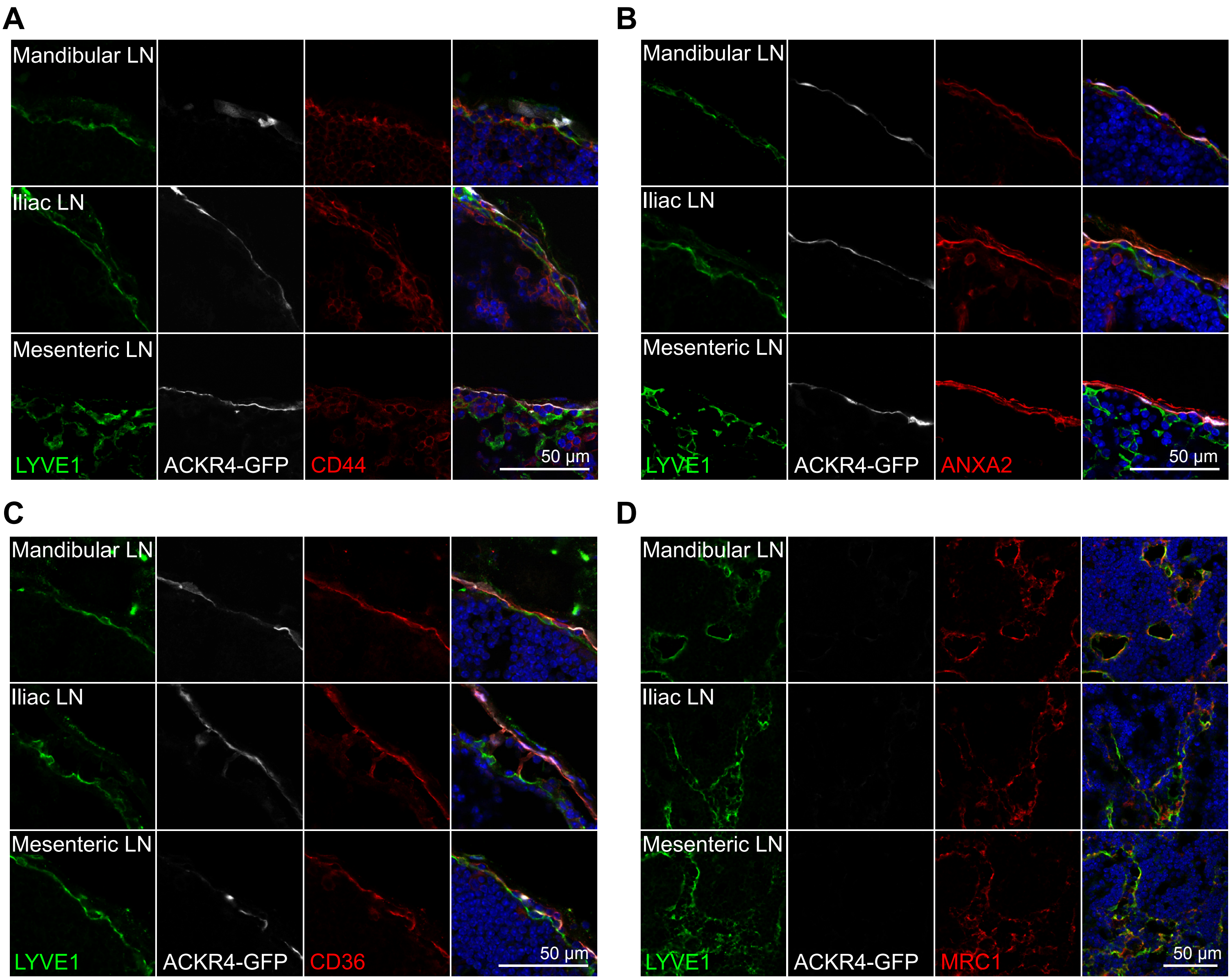

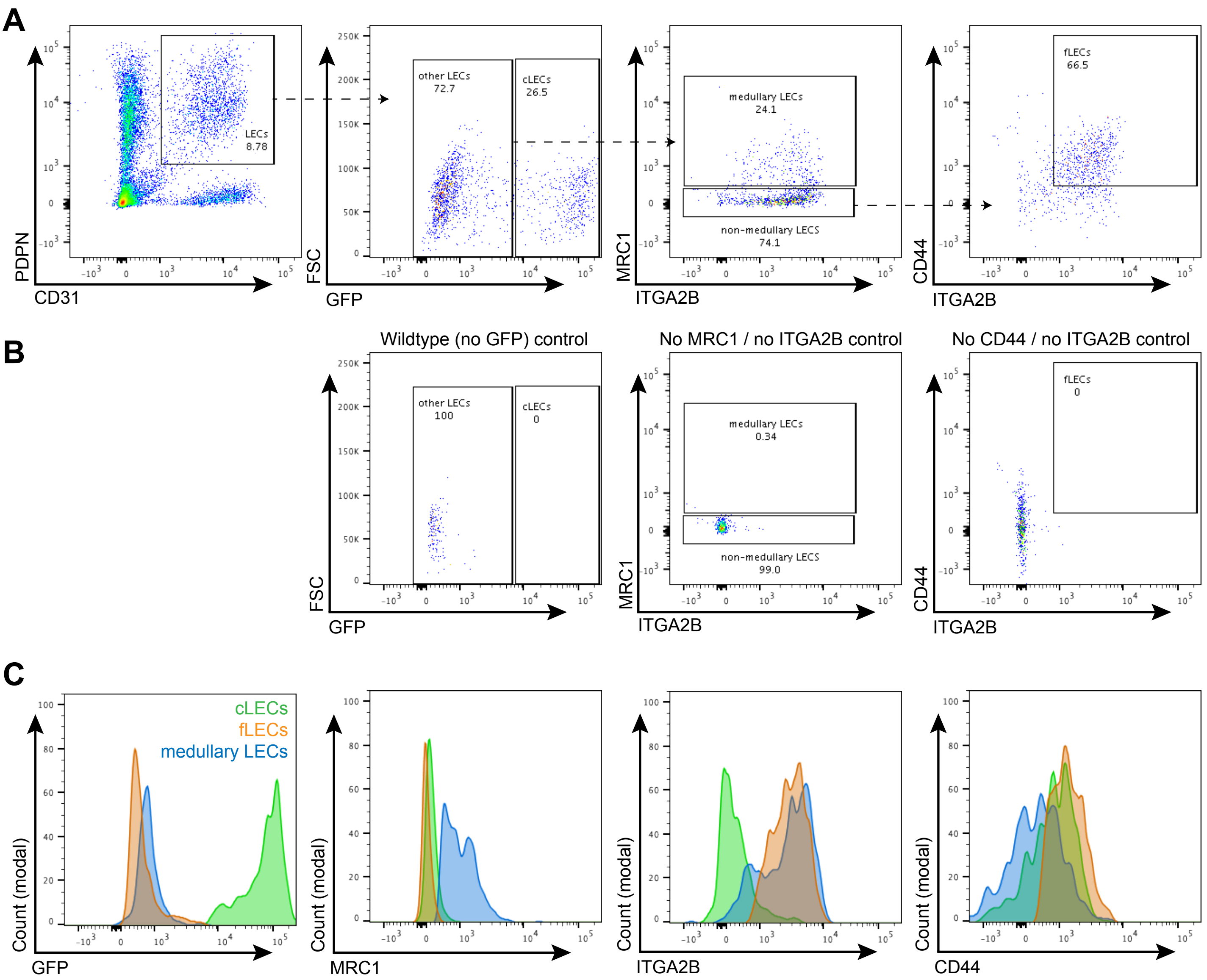

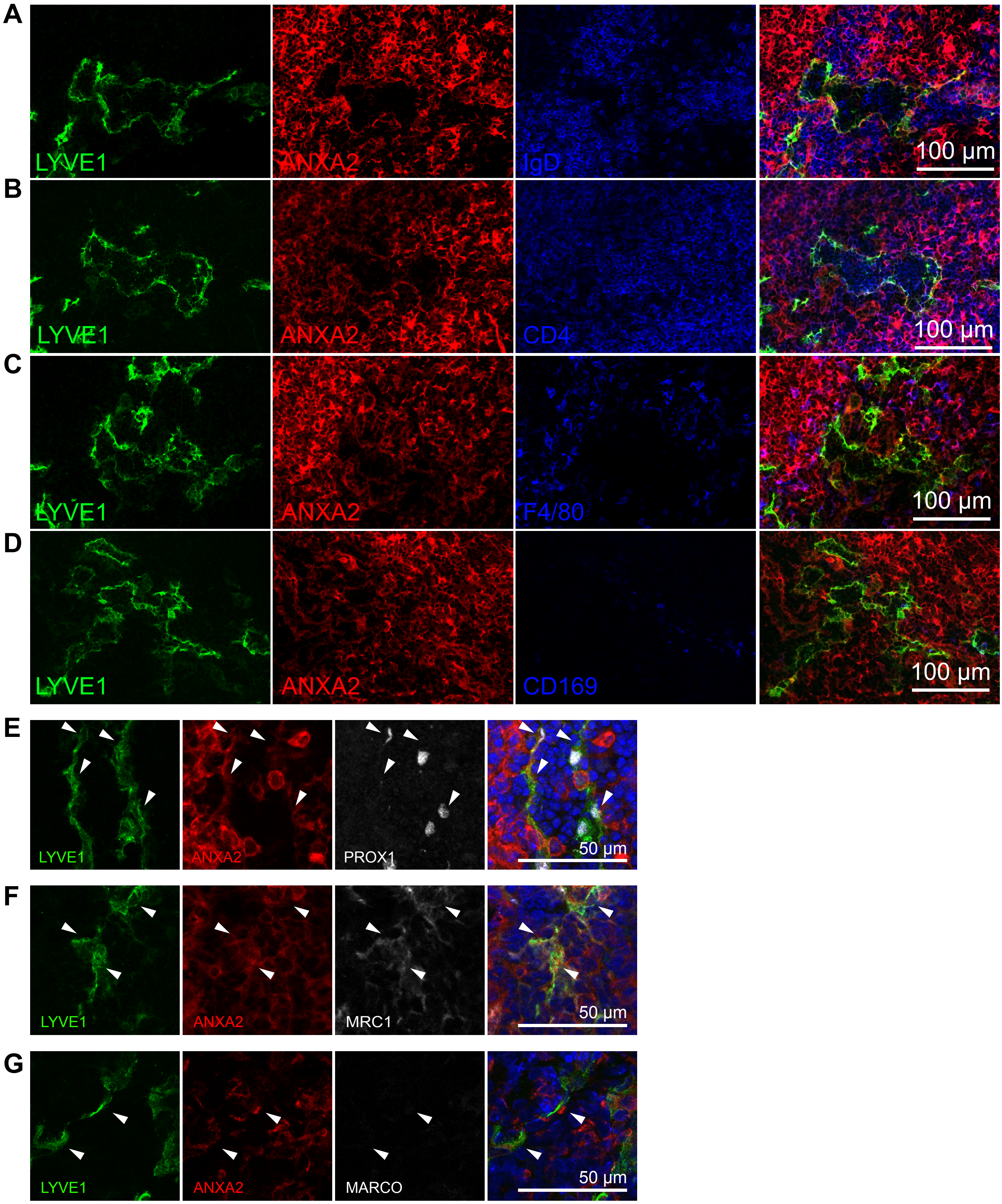
